# Metabolite Fraction Libraries for Quantitative NMR Metabolomics

**DOI:** 10.64898/2025.12.30.696914

**Authors:** Christopher Esselman, Kara Garrison, Leandro Ponce, Ricardo M. Borges, Frank Delaglio, Arthur S. Edison

**Affiliations:** Institute of Bioinformatics, University of Georgia, Athens, Georgia, USA; Department of Biochemistry and Molecular Biology, University of Georgia, Athens, Georgia, USA; College of Engineering, University of Georgia, Athens, Georgia, USA; Complex Carbohydrate Research Center, University of Georgia, Athens, Georgia, USA; Instituto de Pesquisa de Produtos Naturais Walter Mors, Universidade Federal do Rio de Janeiro, Rio de Janeiro, Brazil; Institute for Bioscience and Biotechnology Research, National Institute of Standards and Technology and the University of Maryland, Rockville, Maryland, USA

## Abstract

Nuclear Magnetic Resonance (NMR) has unique strengths in metabolomics studies, particularly in quantifying mixtures and elucidating the structures of unknown molecules. One-dimensional (1D) proton (^1^H) NMR is the most common method; however, spectral overlap is significant, making analysis challenging. We present a new approach that utilizes chromatographically separated fractions from a pooled sample, henceforth called a metabolite fraction library (mFL). We developed an algorithm to extract highly correlated peaks from the mFL, collectively forming a metabolite basis set (mBS). The mBS can be fit to NMR profiling data, enabling comprehensive quantification. Applied to 10 mixtures of 53 metabolites, our approach accurately quantified 50, quantified an impurity and an oxidation product, and described between 91-96% of total spectral intensity. The method is demonstrated using the fungus *Neurospora crassa*, resulting in the identification of 45 metabolites with high confidence, 45 with medium confidence, and accounting for 94% of total spectral intensity.

---

The primary analytical methods for metabolomics research are liquid chromatography-mass spectrometry (LC-MS) and nuclear magnetic resonance spectroscopy (NMR). Recent trends show an increased adoption of LC-MS compared to NMR, although the use of both techniques is growing.^1^ The popularity of LC-MS stems from its high sensitivity, selectivity, and the availability of extensive MS/MS fragmentation databases.^2, 3^

One-dimensional (1D) proton (^1^H) NMR is the most common approach in NMR-based metabolomics due to the high natural abundance of ^1^H, its prevalence in organic compounds, and because ^1^H is the most sensitive to NMR measurement among all the stable isotopes. As such, ^1^H NMR is a near-universal detector of sufficiently concentrated metabolites, and NMR has the further advantage of being inherently quantitative.^4^ Samples are never in contact with the NMR spectrometer, eliminating the need for complicated sample preparation and allowing for few restrictions on sample type. Additionally, since NMR is non-destructive, more experiments can be conducted after initial data acquisition. Finally, the covalent structures of unknown molecules can be determined *de novo* through powerful two-dimensional (2D) NMR correlation methods.^5^

Given the long list of advantages, why is NMR less popular than LC-MS in metabolomics research? We argue that the primary limitation of ^1^H NMR is signal overlap because we do not routinely chromatographically separate our samples. Alternatives exist to 1D ^1^H NMR, most notably various 2D NMR methods.^6–9^ 2D NMR resolves much of the overlap. Still, the experiments take longer to collect, are more challenging to quantify, and are not well-suited for profiling hundreds or thousands of samples.

There are two primary approaches for analyzing 1D ^1^H NMR-based metabolomics data. The chemometric approach uses multivariate analysis to identify peaks in NMR spectra that differentiate two or more groups,^10, 11^ and only important peaks in the study are examined in detail. Related chemometric approaches use statistical correlations across spectra^12^ or ratios of spectra^13^ from different groups to associate peaks into molecules. These methods are powerful, but their performance suffers in regions of overlap.^14^

An alternative to the chemometric approach is direct quantitative analysis of NMR spectra.^4^ These methods utilize reference libraries to fit spectra quantitatively, which works well for targeted studies, but is limited for non-targeted studies. Computational methods are continually improving, so it may eventually be possible to generate comprehensive libraries solely from computed spectra. However, the accuracy of chemical shift prediction is still insufficient for this purpose. More importantly, as the library grows, so do false discoveries, and the overall probability of the correct fit suffers.^15^ One of the most popular quantitative software systems for NMR metabolomics is Chenomx^TM^ NMRSuite.^16^ This commercial software utilizes an extensive database derived from the Human Metabolome Database (HMDB),^17^ which accounts for changes in pH. While Chenomx is powerful for many applications, typical workflows often require subjective, interactive steps, which can lead to results that are not always statistically robust and may be biased.^18^ Bruker has developed another commercial approach to quantifying NMR data as part of their IVDr system.^19, 20^ IVDr works on a limited set of human biofluids and requires particular sample preparation and data collection. The data are quantified and identified in the cloud with proprietary software; therefore, the method has the drawback of not being transparent.

Many academic software tools^4^ have been developed for partial automation of NMR metabolomics quantification. Two tools particularly relevant for the work presented here are Bayesil^21^ and BATMAN.^22^ Both methods use Bayesian statistics with a defined set of NMR reference data that serve as prior knowledge. These approaches are powerful but have some limitations. Bayesil requires NMR spectra to be collected under precise conditions, and quantification requires sample-specific libraries. BATMAN is more general but relies on a defined set of reference spectra, making it most suitable for targeted NMR analysis.

Here, we introduce a novel method for complete quantitative NMR metabolomics that provides a fit-for-purpose approach that simplifies analysis and specifically accounts for the sample type of interest, as an alternative to using pre-existing general reference libraries. The method consists of three main steps, as shown in Fig. 1. Step 1 creates a metabolite fraction library (mFL) using a pooled sample or reference material representative of the biological study.^23^ The sample is chromatographically separated into non-targeted fractions by semi-preparative high-performance liquid chromatography (HPLC). NMR spectra of the fractions are modeled in the time domain using spectral automated NMR decomposition (SAND), resulting in individual time domain signal models that can be Fourier processed in the same way as the measured data to generate peaks.^24^ Step 2 regroups the modeled peaks into known and unknown molecules by performing correlations across fractions. We refer to the collection of regrouped molecules as the *metabolite basis set* (mBS). Step 3 includes the quantitative Bayesian fitting of the mBS to an unfractionated 1D ^1^H NMR profiling spectrum to obtain the concentrations of every mBS element.

**Fig. 1:**
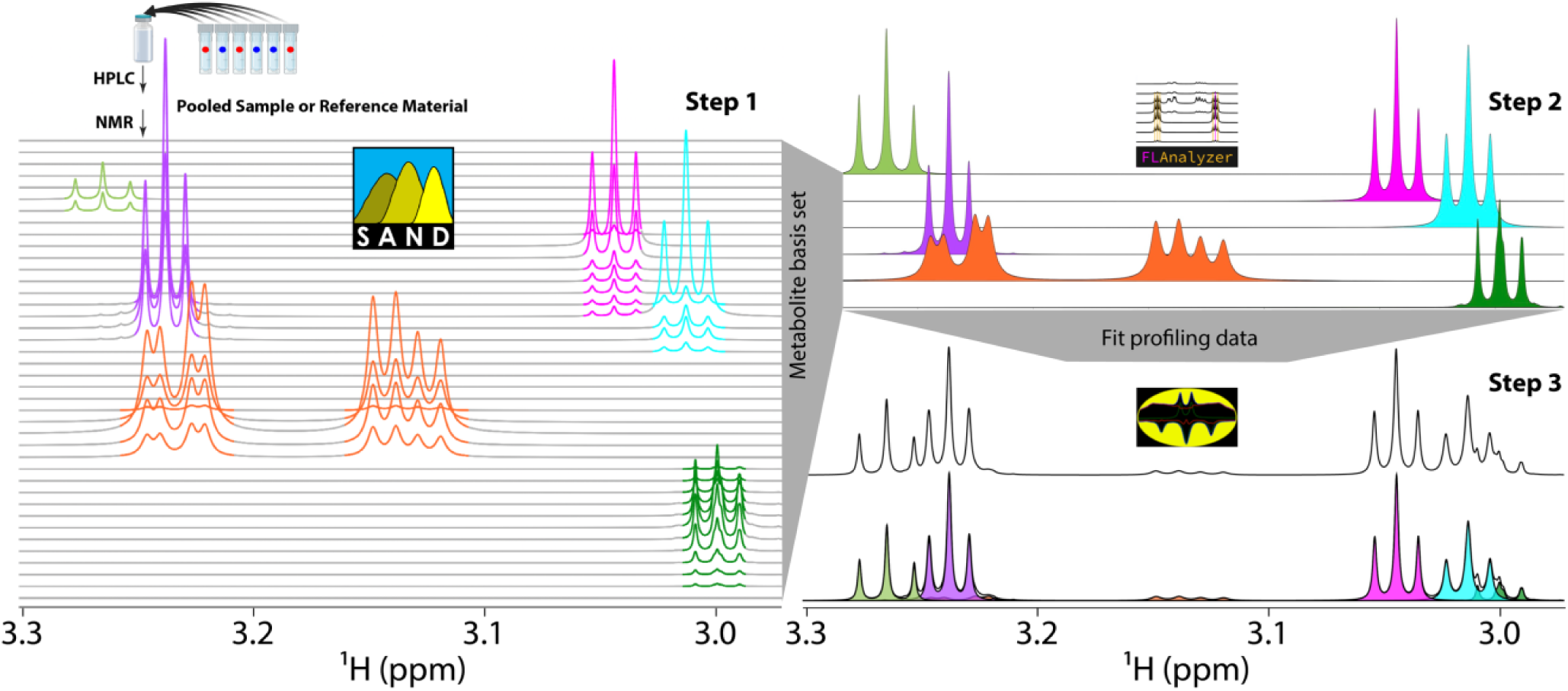
Overview of the method. **Step 1** represents the construction of the metabolite fraction library (mFL), starting with a pooled sample or reference material, followed by HPLC chromatography with untargeted fraction collection, 1D ^1^H NMR spectroscopy of all fractions, and tabular domain creation using SAND. **Step 2** creates the metabolite basis set (mBS) by correlating all peaks in the mFL across eluting fractions using a MATLAB application called FLAnalyzer. The mBS can be database-matched to derive known compound annotations. **Step 3** fits the known and unknown mBS elements into an unfractionated 1D ^1^H NMR spectrum of a mixture from the study. We use BATMAN with the mBS as prior knowledge in a Bayesian fit to the data.

Fractionation in natural products research is routine but less common in metabolomics.^25^ However, some metabolomics studies have effectively incorporated fractionation into compound identification workflows or when integrating NMR with LC-MS.^26–28^ For example, Whiley and coworkers created a “fraction bank” to aid in identifying several urine metabolites using a combination of chromatography, MS, and NMR.^29^

Using 10 different known mixtures of 53 reference samples of metabolites as “ground truth”, we merged fraction library spectra of these reference samples into the mBS, matched them to databases for annotation, and fit them to each of the mixtures. Most of the resulting concentrations had nearly perfect correlation coefficients compared to integrated values of isolated peaks in the mixtures. We demonstrate the utility of our method using a biological sample of the filamentous fungus, *Neurospora crassa*.

## Results

### Metabolite fraction libraries mitigate overlap in NMR spectra

The workflow to generate an mFL is shown in Fig. Supplementary 1, and a region from the mFL derived from *N. crassa* is shown in Fig. 2a. The black trace on the top shows an unfractionated 1D ^1^H NMR spectrum of the same sample, demonstrating the extensive signal overlap typical in conventional 1D ^1^H NMR metabolomics spectra. For example, the doublet at ∼2.48 ppm in fraction ∼35 (box 1) is completely lost in the mixture due to the overlap of the large signals around fraction 110. Similarly, the smaller peaks around 1.9 ppm and 2.3 ppm appear to be in the same molecule but are obscured in the unfractionated sample by two different sets of overlapping peaks (box 2). The unfractionated peak at 1.5 ppm appears distorted, which might be interpreted as poor phasing or shimming, but the fraction library clearly shows different sets of well-defined multiplets in other molecules that cause the distortion (box 3).

**Fig. 2:**
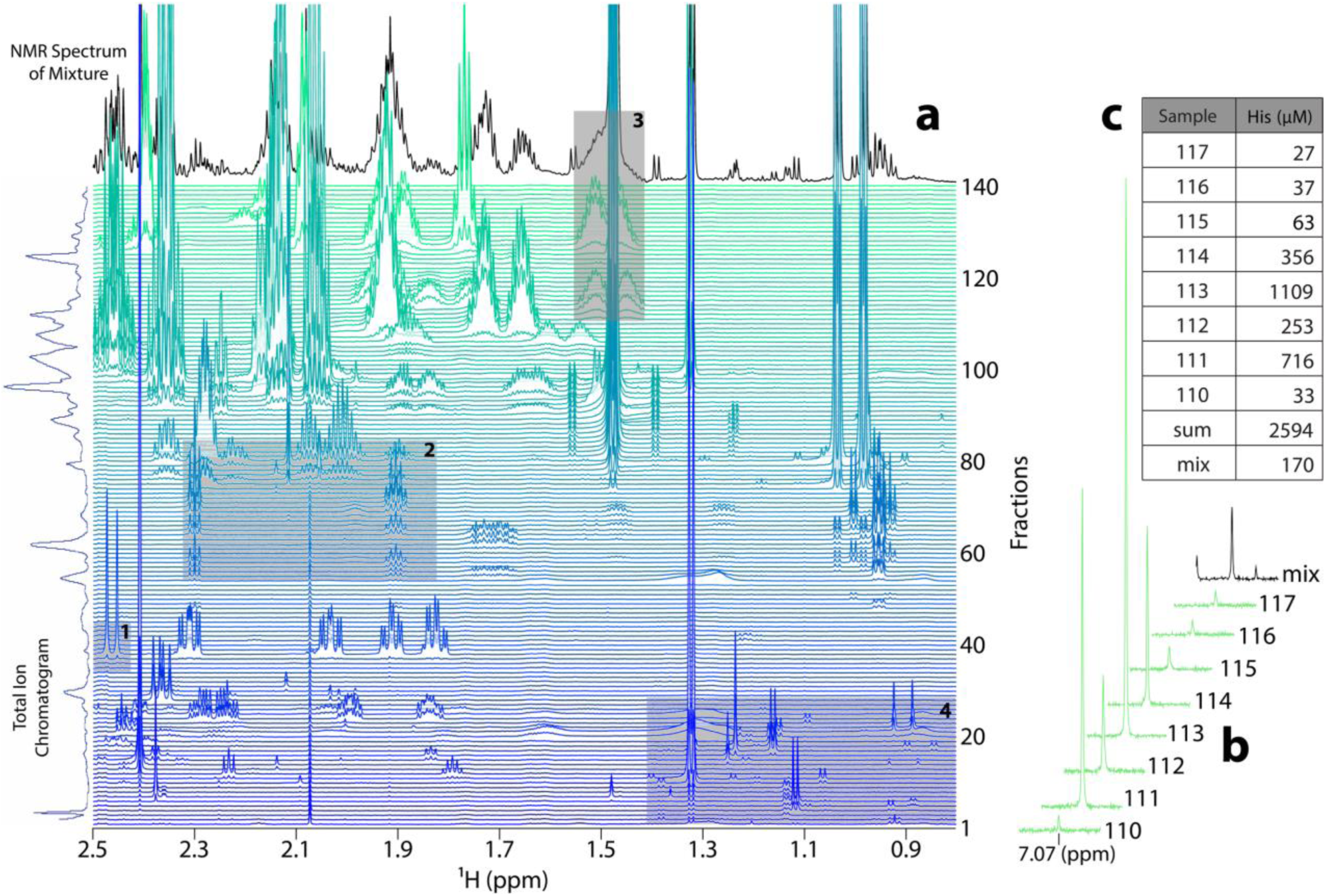
**a**, *N. crassa* metabolite fraction library (mFL), displayed as a spectral series of ^1^H NMR measurements of all fractions. The excerpted spectral region shown includes methyl and aliphatic signals. The black trace at the top is the unfractionated NMR spectrum of the extract used to make the mFL, and the vertical blue trace on the left is the total ion chromatogram from the HPLC separation. The four shaded boxes highlight different regions that are described in the text. **b**, One of the peaks from histidine that resonates at roughly 7.07 ppm across its eluting fractions 110 to 117. The final trace in black is the same histidine peak in the unfractionated mixture. The peak areas are normalized according to the integral of the DSS reference signal in each spectrum. **c**, Table of derived concentrations of all the histidine peaks from **b** reporting mFL histidine concentrations in μM for fractions 110 to 117, the sum of these concentrations (“sum”), and the concentration in the unfractionated mixture (“mix”).

### Metabolite fraction libraries can increase the sensitivity of NMR metabolomics

Several regions of Fig. 2a have small but authentic metabolite peaks in the fraction library (e.g., box 4). Some of these small peaks can be seen in the unfractionated mixture, but others would be impossible to distinguish from noise. This can be surmounted by using auto-injection and repeated HPLC fractionation runs to accumulate several injections into the same fractions. This results in samples with higher concentration, which increases the achievable sensitivity of metabolite detection by NMR.

Fig. 2b shows quantification of this sensitivity increase using histidine peaks that are well-resolved in the unfractionated spectrum. The sample was prepared from 6 mg of *N. crassa* cell mass, and from integration, the concentration of histidine in the mixture is 170 μM. The fraction library used 90 mg of *N. crassa*, and the table insert in Fig. 2c shows the concentrations of histidine in each fraction. The sum of all the fractions is 2594 μM, 15.3 times greater than the unfractionated signal, and essentially the same as the 1:15 ratio of the starting cellular masses. As expected, the NMR signal is proportional to the amount of sample, explaining the benefit of multiple injections.

### SAND time domain modeling compensates for spectral overlap in fraction libraries

Recently, we reported a new method called spectral automated NMR decomposition (SAND) to model NMR data in the time domain, accurately quantifying overlapping peaks without the need for interactive analysis (Fig. Supplementary 2a).^24^ Time domain modeling provides advantages over integration of spectra, because baseline distortions that extend through the spectrum arise from a small number of distorted points at the start of the time domain data, and these can be down-weighted or omitted when fitting the model. The output from SAND is tabular domain data^30^ that includes the frequency, amplitude, decay, and phase of every signal in the modeled data. These parameters can be used to generate synthetic time domain data for each signal, and this synthetic data can be Fourier processed in the same way as the measured data to create model spectra (Fig. Supplementary 2b). Since each signal can be synthesized separately, it is possible to use SAND results to generate spectra of a selected subset of signals, or to subtract selected model signals from the measured data to generate simplified spectra.

### Individual signals identified by SAND can be correlated into molecules

Fig. 3a shows an expansion of the *N. crassa* mFL with the glutamine (Gln) resonances highlighted by purple traces. These traces were generated automatically by an in-house MATLAB application called FLAnalyzer, which identifies peaks that persist over multiple fractions. Correlation analysis is used to identify peak traces whose intensities change in the same way over the fractions as they would if the peaks all arose from the same molecule. Peaks whose traces are correlated above a selected threshold (Pearson’s correlation coefficient *r* > 0.98; Fig. 3b) are grouped into a metabolite basis element (Fig. 3c).

**Fig. 3:**
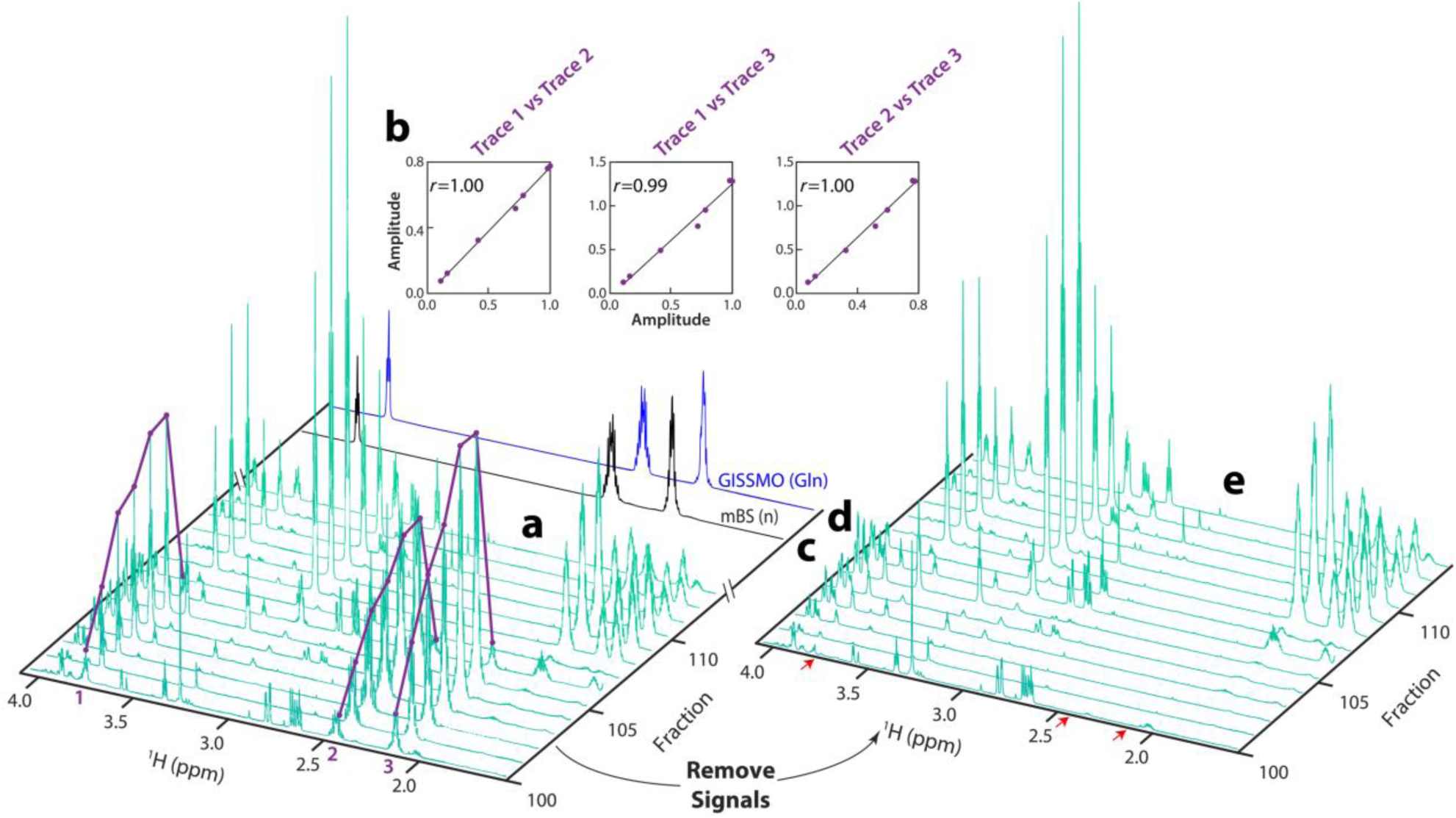
**a**, Region of the mFL following FLAnalyzer tracing of correlated peaks across fractions. The three purple curves highlight SAND peaks at 3.761 ppm (Trace 1), 2.426 ppm (Trace 2), and 2.130 ppm (Trace 3), all of which are highly correlated over a range of fractions. **b**, The values of these correlations across eluting fractions, where the correlation coefficients *r* are all greater than 0.99. **c**, Highly correlated peaks are grouped to create a metabolite basis element (mBS (n), where n indicates that this is the n^th^ element in the mBS. **d**, The mBS elements are then matched to a database to provide annotations, as described in the text. In the case shown here, the three highly correlated signals show a strong match to glutamine (Gln) from the GISSMO library, shown in blue. **e**, Because the mFL is constructed from tabular domain SAND data, the correlated peaks can be removed by subtracting simulated versions of the peaks, resulting in a simplified mFL.

FLAnalyzer selects the highest intensity peak in the mFL dataset and uses it as a driver to find other correlated peaks in the same fractions as the driver peak. Following grouping into a basis element, these peaks can be subtracted from the original mFL, and analysis can be iteratively repeated on the residual (Fig. 3e). This is a unique feature of tabular domain data as produced by SAND, because modeled peaks can be subtracted from the measured data to generate simplified spectra for further analysis. After removing the previous basis set signals, the algorithm starts again with the remaining highest intensity peak in the modified mFL.

### Metabolite basis elements form a nearly complete set of NMR-observable metabolites

The result of FLAnalyzer is the metabolite basis set (mBS), which represents all basis set elements extracted from the mFL (Fig. 4a). In the case of *N. crassa,* we obtained 126 mBS elements. Each element of the mBS can then be matched to a 1D ^1^H NMR database (Table Supplementary 1). We prefer COLMAR1D,^31, 32^ which uses the GISSMO^33^ database, because GISSMO is a spin matrix representation of the data that can generate synthetic spectral data at any NMR field strength.

**Fig. 4:**
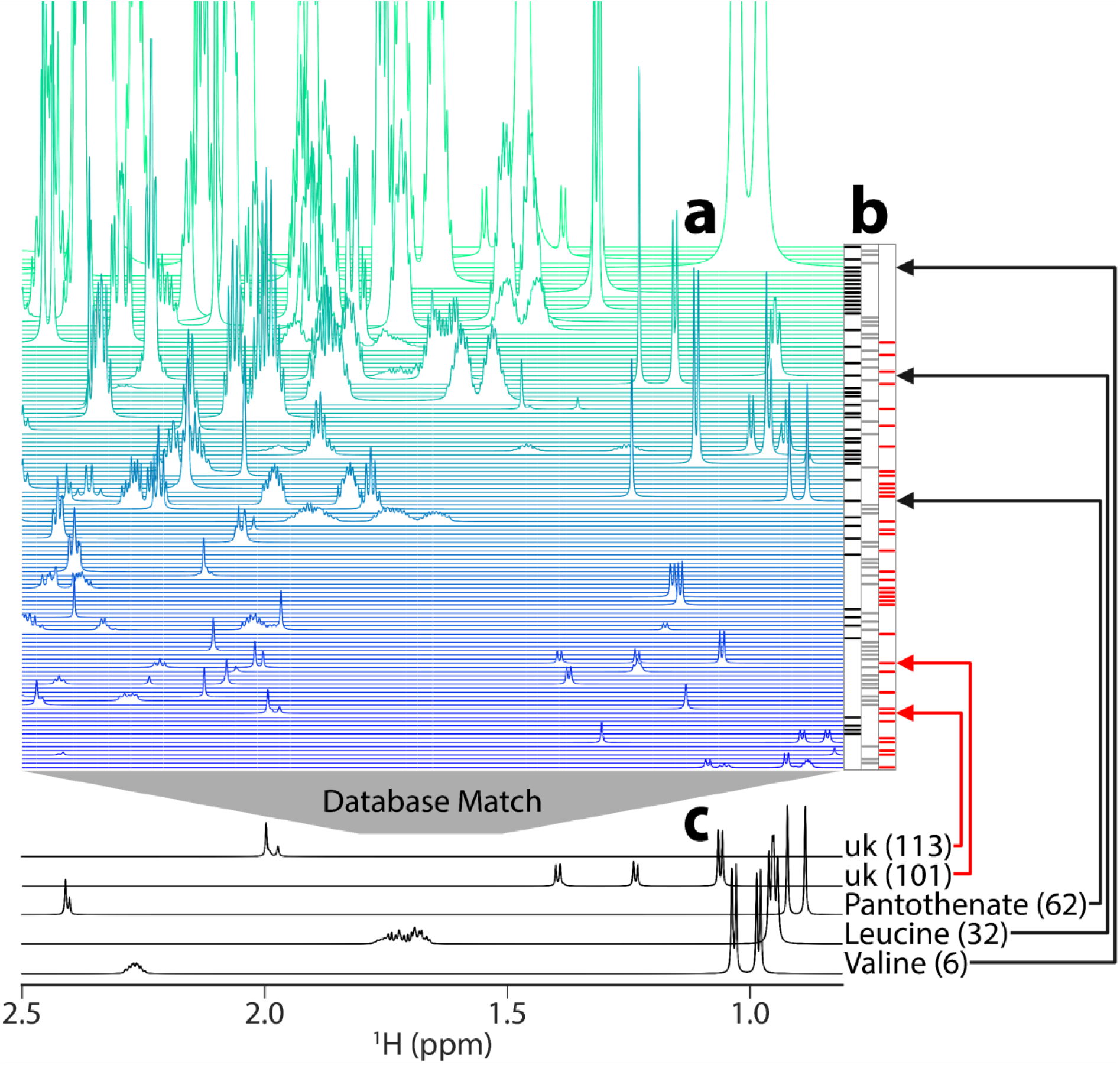
**a**, *N. crassa* metabolite basis set (mBS). This mBS has 126 elements, which correspond to putative metabolites. It is ordered from highest intensity on row 1 (top) to lowest on row 126 (bottom). **b**, Colors represent confidence in database matching with COLMAR1D using the GISSMO reference database. Black indicates database matches with high confidence, grey indicates lower-confidence database matches, and red indicates no database matches. **c**, Five examples of mBS elements over the same chemical shift range as in **a**, with the mBS element numbers in parentheses. Valine, leucine, and pantothenate all led to high confidence matches, and two different unknowns (uk) are shown. Arrows (black for known and red for unknown) point to the corresponding location of each element in the basis set.

### Over 95% of the metabolites are accurately quantified in a ground-truth mixture

To test the workflow outlined in Fig. 1, we made 10 experimental mixtures of 53 metabolites. First, we created an mFL (Step 1, Fig. 1) of the mixture, shown in Fig. Supplementary 3. Next, we used FLAnalyzer to extract the mBS from the mFL (Step 2, Fig. 1). We reconstructed the mBS and found database matches for 50 of the 53 metabolites used to create the mFL (Table Supplementary 2). We discovered that the galactose sample obtained for the study had substantial impurities and very little galactose. Consequently, we could not find galactose in the mBS, but interestingly, we could extract and quantify the impurities from that sample (Fig. Supplementary 4). Similarly, we were unable to match cysteine in the mBS; however, upon inspection of the cysteine sample used in the study, we observed resonances that resembled those of cysteic acid, a product of cysteine oxidation reactions that do not require enzymatic catalysis (Fig. Supplementary 5). These peaks were part of the mBS and were quantified. Lastly, pyruvate could not be identified in the mFL, potentially due to impurities and degradation later observed in the starting sample.

The BATMAN fit for one of the mixtures is shown in Fig. 5. As shown, the residual is relatively flat, with this example mBS fit accounting for 96% of the measured spectral intensity. Over all 10 mixtures, the mBS quantification model accounted for 91-96% of the measured spectral intensities. We then compared these mBS results to conventional quantification by peak integration, using 11 different metabolites having at least one non-overlapping peak in the mixture spectra. Fig. 5 shows the correlation between concentrations derived from conventional integration of peaks in the mixture spectrum vs. the BATMAN-derived concentrations. Most of the *r* values are >= 0.97, with the exceptions being two of the metabolites with significant pH sensitivity, namely ascorbate (*r* = 0.92) and histidine (*r* = 0.79). These lower correlation values can be explained by the fact that such cases can show the largest spectral variation between the compound isolated by fractionation and the compound in its original mixture matrix.

**Fig. 5:**
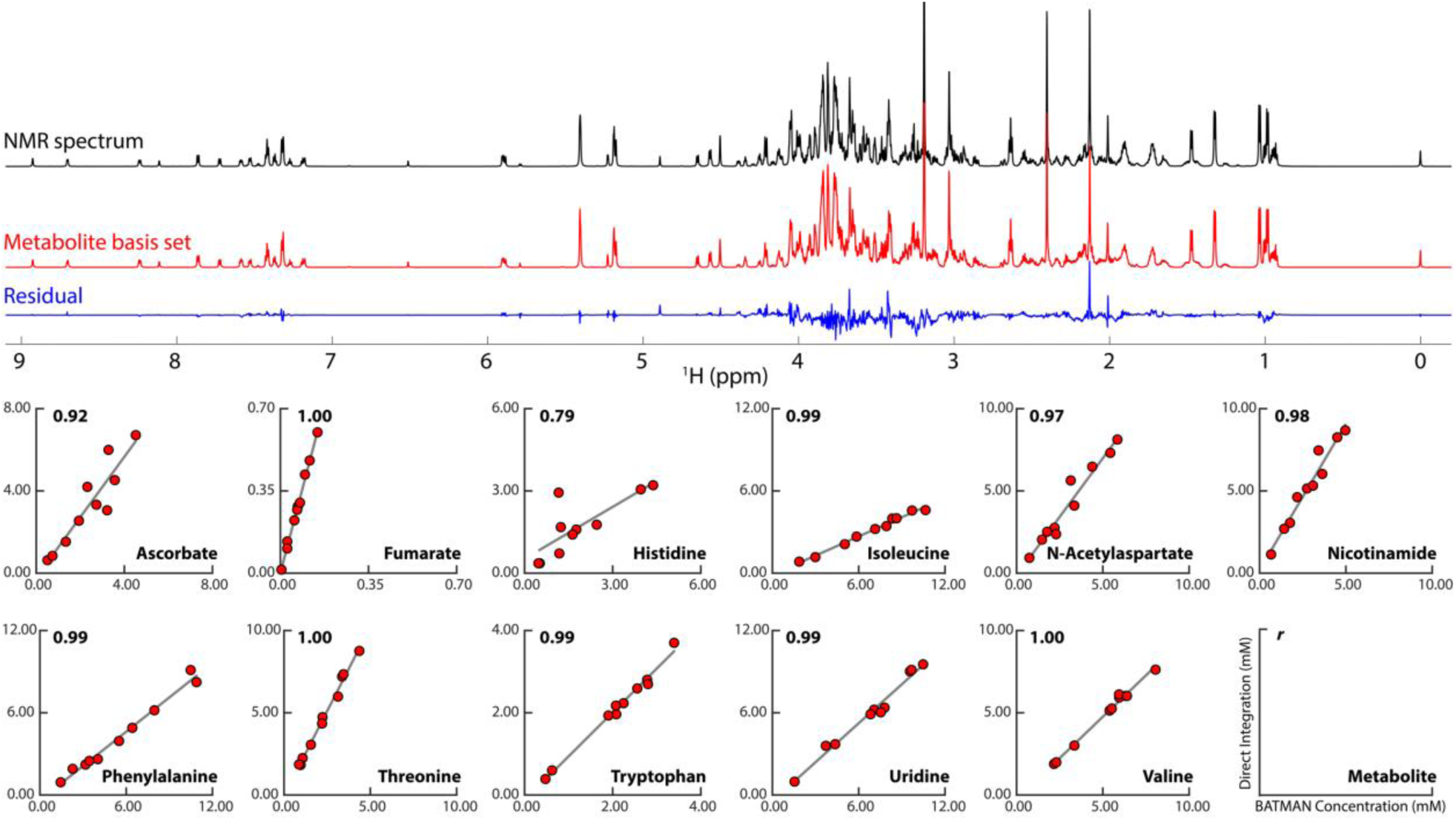
BATMAN fit of the mBS from our ground-truth set of 53 synthetic metabolite solutions. The NMR spectrum at the top (black) is one of ten ground-truth experimental mixtures for this study. The metabolite basis set (mBS) shown in red is the best BATMAN fit to that spectrum from the complete mBS obtained from the ground-truth mFL. The blue trace shows the residuals (wavelet fit) from the Bayesian analysis. For cases where peaks in the mixture spectra could be directly identified and integrated, we compared the results of conventional integration with values extracted using the mBS. The correlation plots at the bottom show BATMAN-derived concentrations vs concentrations derived from numerical integration of peaks in the mixture spectra for 11 metabolites. The numerical values in the upper left of each plot are Pearson’s correlation coefficient *r*, and the name of the metabolite is in the lower right. The integral and BATMAN values were compared to the DSS reference in each spectrum for absolute quantification in mM.

To assess the robustness of BATMAN fitting with the mBS, we held out the first 10 mBS elements and refit the ground-truth mixtures (Fig. Supplementary 6). The percentage of the original spectral intensities quantified fell to 76-88%. We then removed the first 20 mBS elements and refit (Fig. Supplementary 7). The percentage quantified fell again to 47-67%. As each mBS element is identified and removed, the FLAnalyzer approach analyzes the largest remaining peaks in the fraction series, so that the mBS elements are extracted in order of decreasing intensity, with the first element containing the highest intensity peak in the mFL. If 10 and 20 mBS elements are deleted randomly instead, the percentage of intensity quantified improves to 86-95% and 65-87%, respectively.

### Quantitative fit of the N. crassa mBS into an unfractionated sample

To test our method on a biological sample, we used the 126-element *N. crassa* mBS from Fig. 4 to fit the unfractionated 1D ^1^H NMR spectrum of *N. crassa* (Fig. 6). The mBS fit quantified 94% of the total intensity of the spectrum used for fitting, which compares favorably with the ground-truth dataset shown in Fig. 5. Not surprisingly, the sugar region between ∼3 ppm - 4 ppm shows the biggest residuals in both datasets due to high chemical shift similarity between different sugar species. The *N. crassa* fit included both known and unknown mBS elements, providing concentrations for every metabolite reconstructed in the mBS (Table Supplementary 3).

**Fig. 6:**
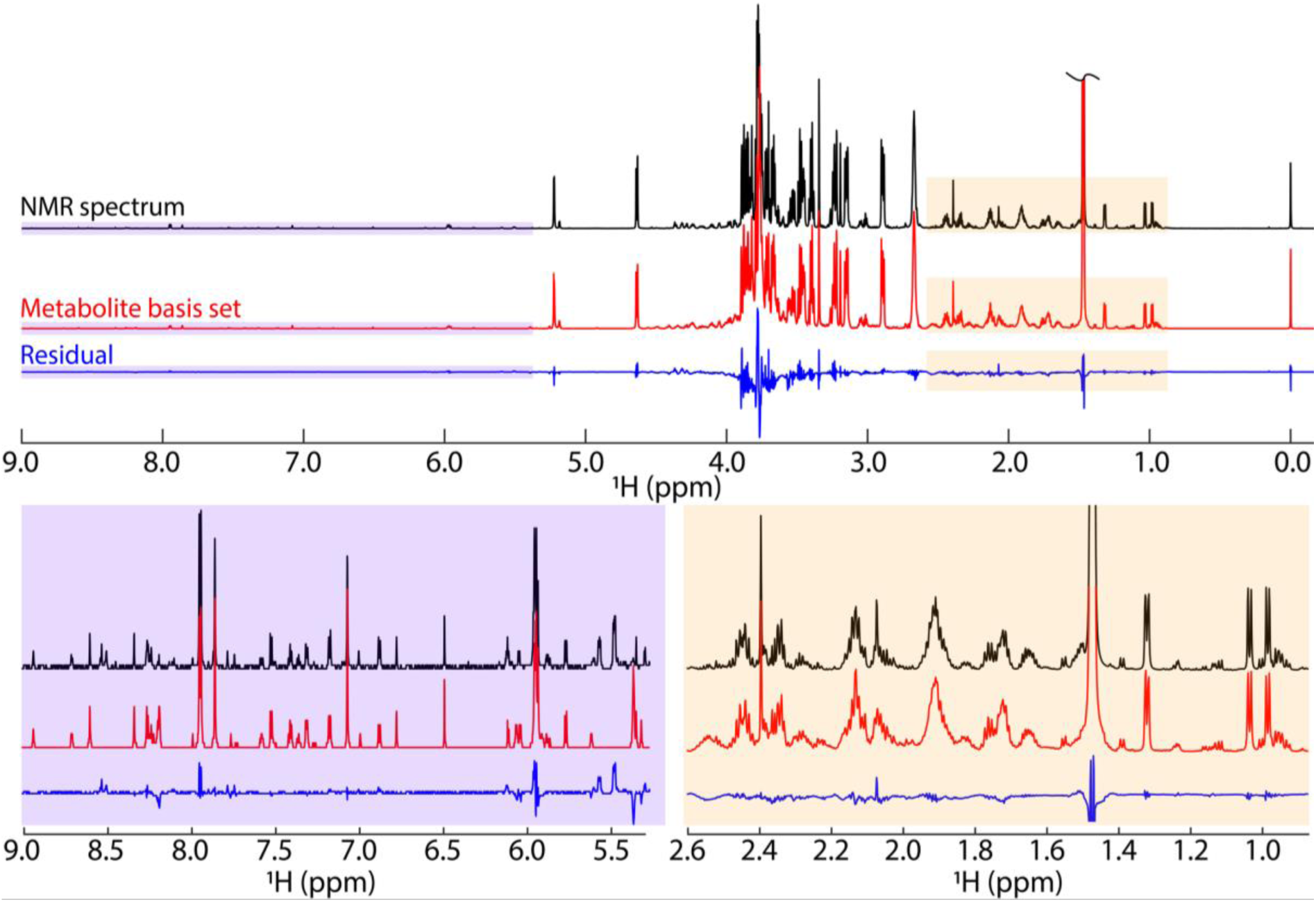
BATMAN fit to the *N. crassa* extract. The NMR spectrum at the top (black) is the original unfractionated material. The metabolite basis set (mBS) shown in red is the best BATMAN fit of the complete mBS obtained from the complete 126 basis elements from Fig. 4. The blue trace shows the residuals (wavelet fit) from the Bayesian analysis. Two expansions to show greater detail are provided.

Next, we removed the first 10 and 20 elements from the mBS and refit the unfractionated spectrum. The percentages of intensity quantified fell to 25% and 18%, respectively, a much steeper drop compared to holdouts in the ground-truth mixtures.

### Quantitative fitting of each mBS across experimental conditions

To investigate the specificity of each mBS to its corresponding experimental spectrum, we fit the ground-truth mixtures with the *N. crassa* mBS and vice versa (Fig. Supplementary 8). The *N. crassa* mBS fit quantified 73-88% of the intensity from the ground-truth mixture spectra, which is lower than the ground-truth mBS fit of the same mixes. Similarly, the ground-truth mBS fit only quantified 83% of the unfractionated 1D ^1^H NMR spectral intensities of *N. crassa*.

## Discussion

We have shown that 1D ^1^H NMR spectra of metabolomics mixtures can be quantitatively fit using metabolite fraction libraries and derived metabolite basis sets in place of a database. This fills a major gap in quantitative NMR metabolomics, which has previously been unable to account for unknown molecules in mixtures effectively. The fractionation approach also enables the identification of features that would be difficult or impossible to find in a heavily overlapped mixture spectrum. Furthermore, conventional quantitative NMR approaches rely on databases, which can work well for targeted studies but fall short for non-targeted studies.

The key step in our new method is creating the metabolite basis set (mBS) from the metabolite fraction library (mFL). The mBS is constructed from mFL spectra separated by HPLC in the chromatographic dimension and peaks accurately decomposed by SAND in the NMR dimension. These steps allow for high correlation thresholds to reconstruct the mBS elements using methods similar to STOCSY.^12^ The relative lack of overlap in mFLs constructed with SAND-processed NMR spectra enables us to use starting thresholds of about 0.98 for correlation constants. Thus, we are confident that mBS elements represent true metabolites in the sample.

Even with the chromatographic and spectral resolution afforded by mFLs, spectral overlap can still interfere with analysis, so FLAnalyzer does include options for interactive adjustment of the automatically determined mBS elements. Signals that are deemed wrongly included or missing in an mBS element can be interactively removed or added in the workflow. However, this step adds significant operator time to the process, as well as the corresponding subjectivity of an individual operator’s interpretation. We are working to refine the fully automated FLAnalyzer analysis to make it both more efficient and less subject to potential bias.

Because the overall sensitivity of an mFL depends on the amount of sample fractionated, it is possible to lower the concentration threshold for NMR metabolomics studies. However, it is not always best to simply add more material. With our system (Agilent 1260 Infinity HPLC), we need to concentrate fractions in a rotary evaporator about every two injections. This adds considerable time and effort to what is otherwise a simple process of creating a fraction library. However, there may be occasions when that extra effort is worthwhile.

The default process for creating the mBS using FLAnalyzer begins with the highest intensity peak in the mFL and proceeds down to the lowest intensity peaks. When the fractions have been highly concentrated, FLAnalyzer can find extremely small peaks for mBS creation. This is primarily because we can subtract the tabular domain SAND peaks as they are correlated into mBS elements. This is a major advantage of SAND processing, because the subtractions are free from distortion and quantitative. As the larger peaks are removed, the smaller peaks become easier to analyze, but this can result in many basis set elements.

BATMAN allows for fine-tuning chemical shifts of multiplets; however, for more pH-sensitive molecules, such as histidine, large shifts can still affect quantitation. This issue can become exacerbated when the spectra used for fitting differ from one another, as seen in the ground-truth mixtures in this study, which have substantial changes in metabolite concentration from sample to sample. Computational techniques have been published^34–37^ to align peaks with chemical shift variation, which may improve quantification in our method. Similarly, SAND tabular domain data may offer opportunities to develop new alignment techniques to assist in this workflow.

## Online Methods

### Data availability

Code and processed data can be found on GitHub: (https://github.com/edisonomics/FLAnalyzer). Raw NMR data can be found on the NAN resource connector.

### Growing Neurospora crassa

Frozen *N. crassa* (NCU06022) stock ordered from the Fungal Genetics Stock Center was used to inoculate Vogel’s solid media growth slants. The slants were incubated in darkness at 30°C for 2 days, then transferred under a benchtop lamp at 25°C for an additional 2 days. The spores were then harvested by washing the slants with 30 mL of autoclaved ddH_2_O and filtering over a cheesecloth.

Polystyrene 14 mL round bottom tubes were prepared with 5 mL of liquid Bird’s Media,^38^ and spores were inoculated at 1 x 10^6^ spores/mL. The tubes were incubated in darkness at 30 °C at 180 rpm overnight. The resulting biomass was vacuum-filtered and washed with 15 mL of autoclaved ddH_2_O. The biomass from every two round-bottom tubes was combined into a 2 mL cryovial. After all the biomass was filtered, the cryovials were flash-frozen with liquid nitrogen and stored in a -80°C freezer.

### Extraction of N. crassa

Ten 2 mL cryovials of biomass were lyophilized for 24 hrs at room temperature, resulting in approximately 90 mg of dried biomass. Five 1.0 mm zirconia beads were added to each tube, and they were bead-beaten for 90 s at 1800 rpm with dry ice using a FastPrep-96 system. 300 µL LC/MS grade 80:20 MeOH/H_2_O was pipetted into each tube and vortexed for 20 seconds. The tubes were then centrifuged at 14,000 rpm at 4 °C for 1 hour. The supernatant was transferred into a single tube and dried using a Centrivap at room temperature. After storage in a -80°C freezer, the resin was reconstituted in 300 µL LC/MS grade 80:20 MeOH/H_2_O and transferred to a 2 mL HPLC Vial with an insert. With the cap on, the vial was spun using a Centrivap without vacuum during the HPLC setup to push any undissolved particulate to the bottom of the vial.

Another cryovial of approximately 6 mg of dried biomass underwent the same extraction procedure and was used as the mixture for fitting.

### HPLC Fractionation

The fraction library was produced by three 100 µL injections using an Agilent 1260 Infinity HPLC with XBridge BEH Amide OBD Prep Column, 130 Å, 5 µm, 10 x 250 mm HILIC column at 25°C. Full scan data were collected using an Agilent Infinity Lab Single Quadrupole MSD in positive ion mode (130-1250 Da). A 38 min linear gradient of 0.1% formic acid in H_2_O (A) and 0.1% formic acid in ACN (B) was used for the fractionation. From 0-20 min, a linear gradient of 5% to 30% A was used, followed by a linear gradient of 30% to 50% A from 20-30 min, all at a flow rate of 3.5 mL/min.

From 30-35 min, a linear gradient of 50% to 65% A was used, followed by an isocratic hold from 35-38 min, both at a flow rate of 2 mL/min. A post-time of 8 min was set to allow the system to equilibrate to the initial condition of 5% A before further injections. Between 4.3 and 30 min, 140 equally spaced fractions were collected, approximately 11 s per fraction. Fractionation was done over identical fraction vials for all three injections. Between injections 2 and 3 and after injection 3, the vials were dried using a Centrivap. The dried fraction vials were stored at -80 °C before NMR data collection.

### NMR Data Acquisition and Processing

The fractions and mixture were reconstituted in 55 μL D_2_O buffer and transferred to 1.7 mm Bruker SampleJet tubes. Solvent blanks were placed in positions 1, 96, 97, and 144 amongst the fractions. NMR data were collected using a Bruker Avance Neo console on an Oxford 800 MHz magnet with a 1.7 mm TCI cryoprobe and a cooled SampleJet sample changer. One-dimensional NMR data were acquired at 298K using a “noesypr1d” pulse sequence, and 32,768 points were collected with 8 dummy scans and 64 scans for each sample. The data were automatically updated to the Network for Advanced NMR (NAN) resource connector and NMRbox for processing.

### Data Processing and Analysis

The following processing and analysis were performed on NMRbox.^39^

#### Reference Deconvolution

Since the SAND modeling procedure uses exponential decay models for the time domain signals, adjusting the data by preprocessing steps to compensate for non-ideal aspects of the experimental lineshapes is beneficial. Our application uses reference deconvolution for this purpose. Reference deconvolution assumes that the entire spectrum has been distorted by a linear, frequency-invariant broadening operator 𝐺(𝑓). This operator acts equally across the entire spectrum; therefore, the distortion correction can be treated as a single operation.^40, 41^ Because of this, a reference signal (DSS singlet peak) 𝑆_𝑟𝑒𝑓_ (𝑓) is assumed to have the same distortion as the rest of the spectrum, and would be composed as: 𝑆_𝑟𝑒𝑓_ (𝑓) = 𝑆_𝑖𝑑𝑒𝑎𝑙_ (𝑓) ∗ 𝐺(𝑓), where 𝑆_𝑖𝑑𝑒𝑎𝑙_ (𝑓) is a perfect ideal peak with no experimental distortions. 𝐺(𝑓) represents all the instrument and acquisition distortions, like poor shimming or local 𝐵_0_ field inhomogeneities.^40, 42^

In a similar fashion, applying this assumption to the whole experimental spectrum’s signal: 𝑆_𝑒𝑥𝑝_ (𝑓) = 𝑆_𝑡𝑟𝑢𝑒_ (𝑓) ∗ 𝐺(𝑓), where 𝑆_𝑡𝑟𝑢𝑒_ (𝑓) is the signal of the whole spectrum without any of the instrumental imperfections.^40, 43^

By the convolution theorem, the frequency domain relationship between 𝑆_𝑡𝑟𝑢𝑒_ (𝑓) and 𝐺(𝑓) can be addressed in the time domain with a point-wise operation, allowing us to deconvolve the desired signal in the following manner: 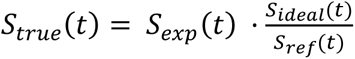. And the desired spectrum corresponds to the Fourier transform of 𝑆_𝑡𝑟𝑢𝑒_ (𝑡). The time domain operation is equivalent to the deconvolution in the frequency domain and is essentially a correction applied at every point of the FID.^40, 41, 44^

Practically, there are important details to consider when implementing reference deconvolution. Namely, the division of 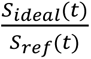 can result in over-amplification of the final points in the time-domain data, which in turn results in sinc wiggle artifacts in the frequency spectrum. Our implementation stabilizes results by applying both an additional exponential window and a gentle trapezoidal apodization to reduce truncation before the final Fast Fourier Transform.

While choosing the parameters of the ideal signal 𝑆_𝑖𝑑𝑒𝑎𝑙_ (𝑡), it is important to note that if a target line width 𝑤 is smaller (in Hz) than the natural linewidth of the experimental spectrum, peaks across the sample will acquire negative artifacts that distort the expected result.^40^ This frequency-independent correction can be used as a preprocessing step to improve the spectra. If a true singlet is unavailable, a multiplet can serve as the reference provided its predictable time-domain zeros are handled (e.g., by interpolation) to yield a stable correction function.^42^

The *N. crassa* data were not reference deconvoluted. However, the ground-truth data were reference deconvoluted with a line width of 1 Hz, a line broadening of 1.5 Hz, and trapezoidal apodization of 75%. A trapezoidal apodization of 75% means the signal tapering begins at 75% of the FID length.

#### SAND

Both fraction libraries and reference mixtures were autophased and processed using NMRPipe 1D batch processing facilities. A script for performing the batch processing can be found at protocols.io (doi: https://dx.doi.org/10.17504/protocols.io.kqdg3xen1g25/v1). In short, the batch processing script applies an automatic zero-order phase correction, a 0.3 Hz exponential line broadening, and automatic first-order baseline correction. The script scales the data so that the maximum value over the whole series is 100. Finally, the script references the DSS peak in each spectrum to 0.0 ppm. The *N. crassa* mixture used for fitting was processed using the same procedure, but since the *N. crassa* spectra exhibited more truncation artifacts and baseline distortion compared to the metabolite reference spectra, a 1.0 Hz exponential line broadening and automatic fourth-order baseline correction were applied. The *N. crassa* fraction library and mixture were then modeled by SAND over the range of 9.0 ppm to -0.5 ppm.

#### SAND frequency reconstruction

SAND models each time domain signal as an exponentially decaying complex sinusoid with amplitude 𝐴_𝑘_, frequency 𝑓_𝑘_, decay 𝜆_𝑘_, and phase 𝜙_𝑘_. Given this signal model, SAND determines the optimal number of signals needed to describe the measured time domain data and determines their parameters. For each signal 𝑘, SAND constructs a time domain model of the form:

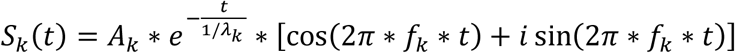

Once these parameters are tabulated, SAND generates model time domain signals using the same digital resolution and parameters as the measured data. The simulated time domain data are saved in the same format as the measured time domain data, so that the corresponding synthetic spectra can be generated by Fourier processing the time domain model in the same way as the measured data. The individual synthetic peak spectra can be summed to yield a composite spectrum that mimics the measured spectrum, and it is possible to delete or subtract unwanted signals (e.g., solvent) before further analysis. As a typical post-processing step to allow point-by-point comparison and analysis, all spectra, both measured and synthetic, are interpolated to generate results with identical size and chemical shift range.

#### FLAnalyzer

After SAND analysis, the output SAND signal tables in comma-separated value format (CSV files) are consolidated into a single directory for correlation analysis. We created a MATLAB application, “FLAnalyzer,” to use the signal tables as input to generate a basis set of metabolites semi-automatically. The application instructions can be found at protocols.io (doi: https://dx.doi.org/10.17504/protocols.io.yxmvmm4jov3p/v1).

#### BATMAN

After making basis sets, the necessary files for quantitatively fitting the mixtures via BATMAN were created. The instructions for generating the BATMAN files can be found at protocols.io (doi: https://dx.doi.org/10.17504/protocols.io.8epv5owmng1b/v1). Several iterations of optimization were conducted to fit the mixtures, as outlined in Hao *et al*. (2014).

#### Calculating Percent Quantified

The percentage quantified by each BATMAN fit was calculated using the equation below.

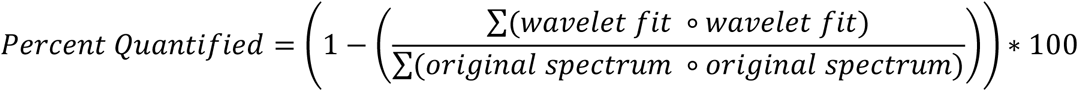

The multiplication shown in the equation represents the element-wise multiplication of each element in the wavelet fit and original spectrum vectors.

#### Database Matching

Peak picking was performed on each basis set element, and the chemical shifts of each peak were used as input for COLMAR 1D Query.^45^ The top database matches were then visually validated as a match by comparing their GISSMO^33^ or BMRB^46^ spectra to the spectra of the basis set elements.

#### Reference Samples and Spectra of Metabolites

Methods for generating samples and spectral data of metabolites used as ground-truth can be found in the supplementary section.

## Author Contributions

Chris Esselman – Produced mFL for ground-truth samples and *N. crassa.* Collected and processed all NMR data. Assisted in the SAND of ground-truth and *N. crassa* mFLs. Wrote all the code of FLAnalyzer and the utilization of mBS with BATMAN. Created linked protocols in protocols.io. Produced all figures. Wrote the manuscript with Edison.

Kara Garrison – Made metabolite ground-truth solutions and mixtures. Assisted in the production and collection of the ground-truth mFL. Assisted in the SAND of ground-truth mFL. Contributed to the methods section for the ground-truth fraction library.

Leandro Ponce – Wrote code for reference deconvolution and sand frequency reconstruction. Assisted in the SAND of *N. crassa* mFL. Wrote the methods section for reference deconvolution and sand frequency reconstruction.

Ricardo M Borges – Developed the idea of correlating peaks as they elute over fractions. Assisted in the HPLC fractionation method and gradient

Frank Delaglio – provided software and guidance for reference deconvolution and SAND analysis to reconstruct metabolite basis sets and edited the manuscript and figures.

Arthur S. Edison – Supervised and conceived of the project. Wrote the manuscript with Esselman and edited figures.

## Acknowledgements

We thank Drs. Brianna Garcia and Goncalo Gouveia for foundational work on fraction libraries. Alexis Molina kindly created *N. crassa* spore suspension stocks. Matt Jacoby assisted in the development of the *N. crassa* extraction protocol. Conrad Epps and Laura Morris beta tested FLAnalyzer. Stephanann Costello provided feedback on the written manuscript. John Grimes helped with NMR automation. Tim Ebbels helped with BATMAN. Óscar Millet provided helpful feedback. This work was supported by NIH 5R35GM148240-03 and the NSF Network for Advanced NMR (1946970).

## NIST Disclaimer

Certain commercial equipment, instruments, and materials are identified in this presentation in order to specify the experimental procedure. Such identification does not imply recommendation or endorsement by the National Institute of Standards and Technology, nor does it imply that the material or equipment identified is necessarily the best available for the purpose.

## Supplement

**Figure Supplementary 1:**
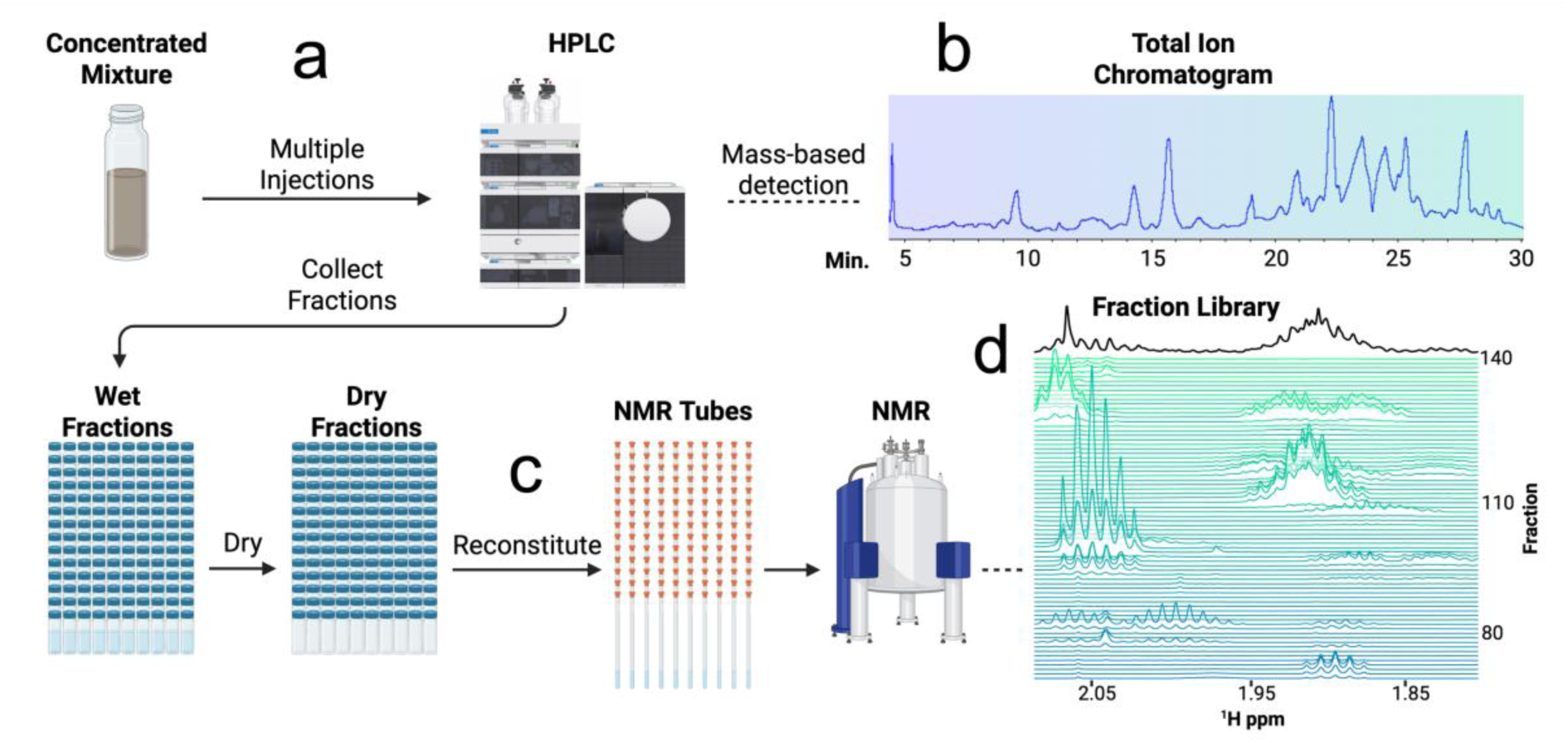
Experimental workflow of creating a metabolite Fraction Library (mFL). **a**, A concentrated biological mixture is injected multiple times into an HPLC using the same fraction vials for each injection. **b**, If a mass spectrometer is coupled to the HPLC, a total ion chromatogram can be collected during fractionation. **c**, The fractions are then dried using a vacuum concentrator and reconstituted into NMR tubes. **d**, One-dimensional (1D) proton (^1^H) NMR is collected for each fraction.

**Figure Supplementary 2:**
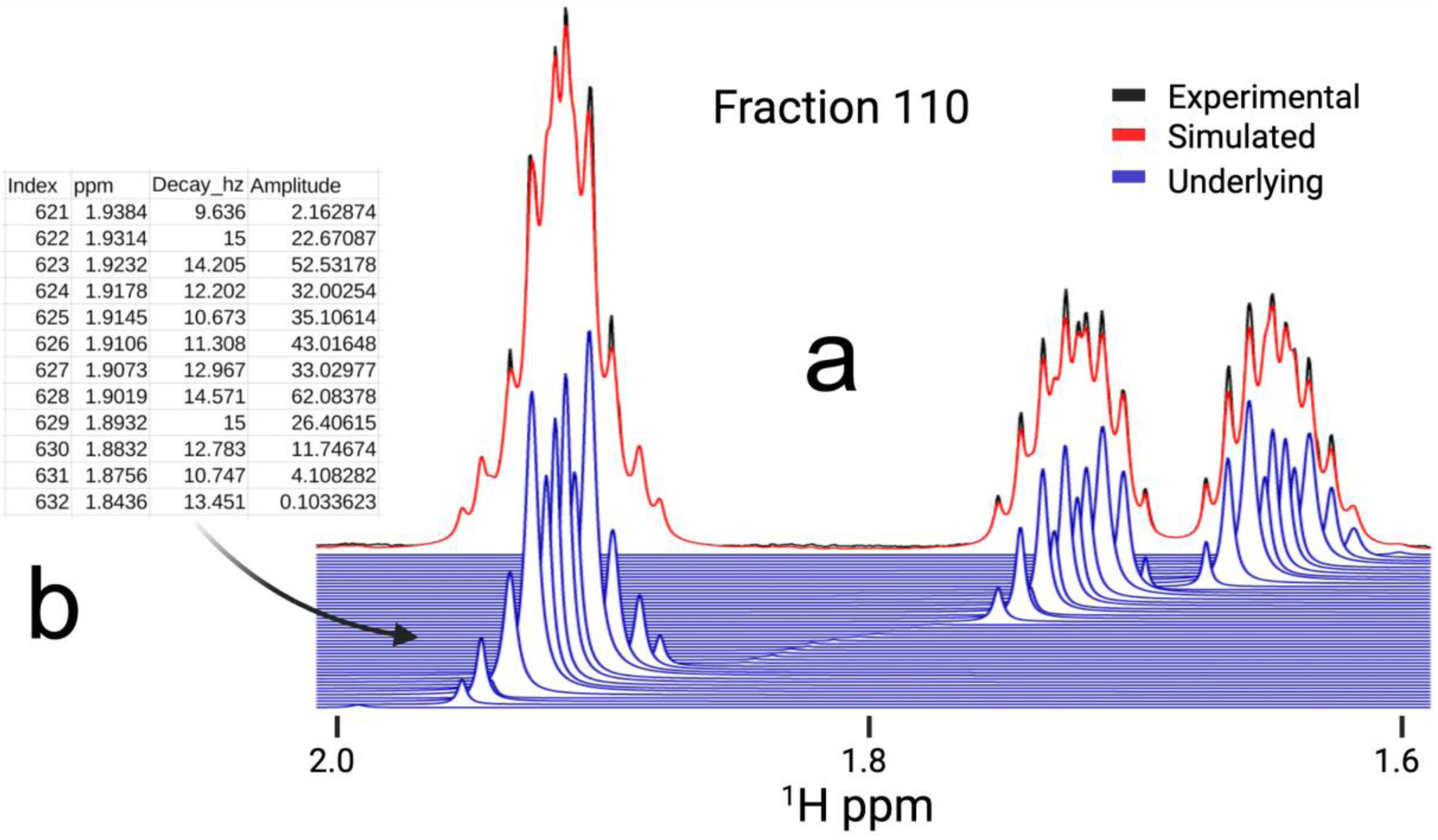
Time domain modeling of NMR data via spectral automated NMR decomposition (SAND). **a**, The black spectrum is the original Fourier transformed (FT) data. The blue peaks are the frequency reconstructed underlying Lorentzian peaks time domain modeled by SAND. The red spectrum is the summation of the blue Lorentzian peaks. **b**, The SAND tabular domain data includes the frequency, decay, amplitude, and phase of every peak in the modeled spectrum. The phase is usually adjusted to zero before SAND.

**Figure Supplementary 3:**
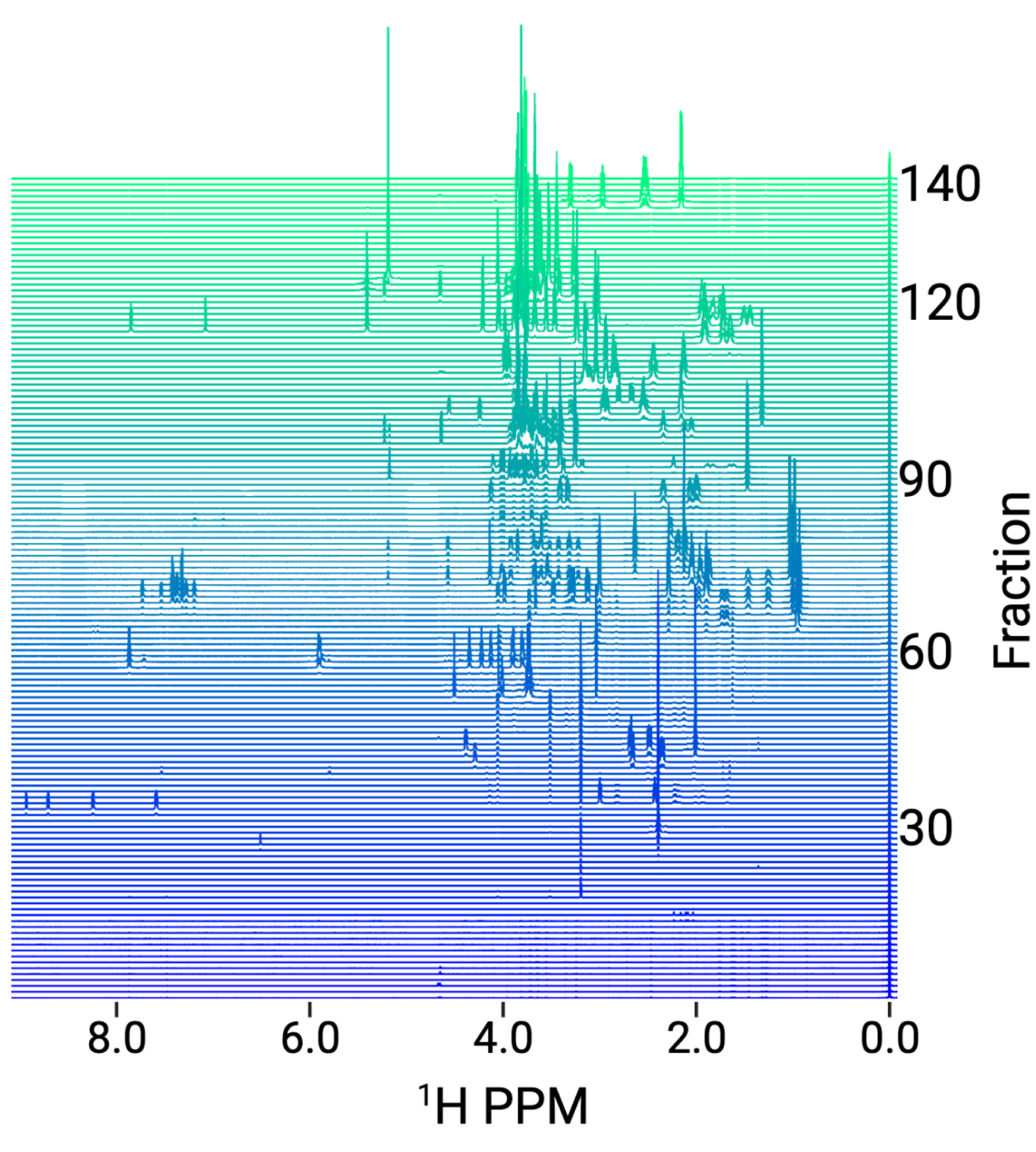
Metabolite fraction library (mFL) of ground-truth mixture.

**Figure Supplementary 4:**
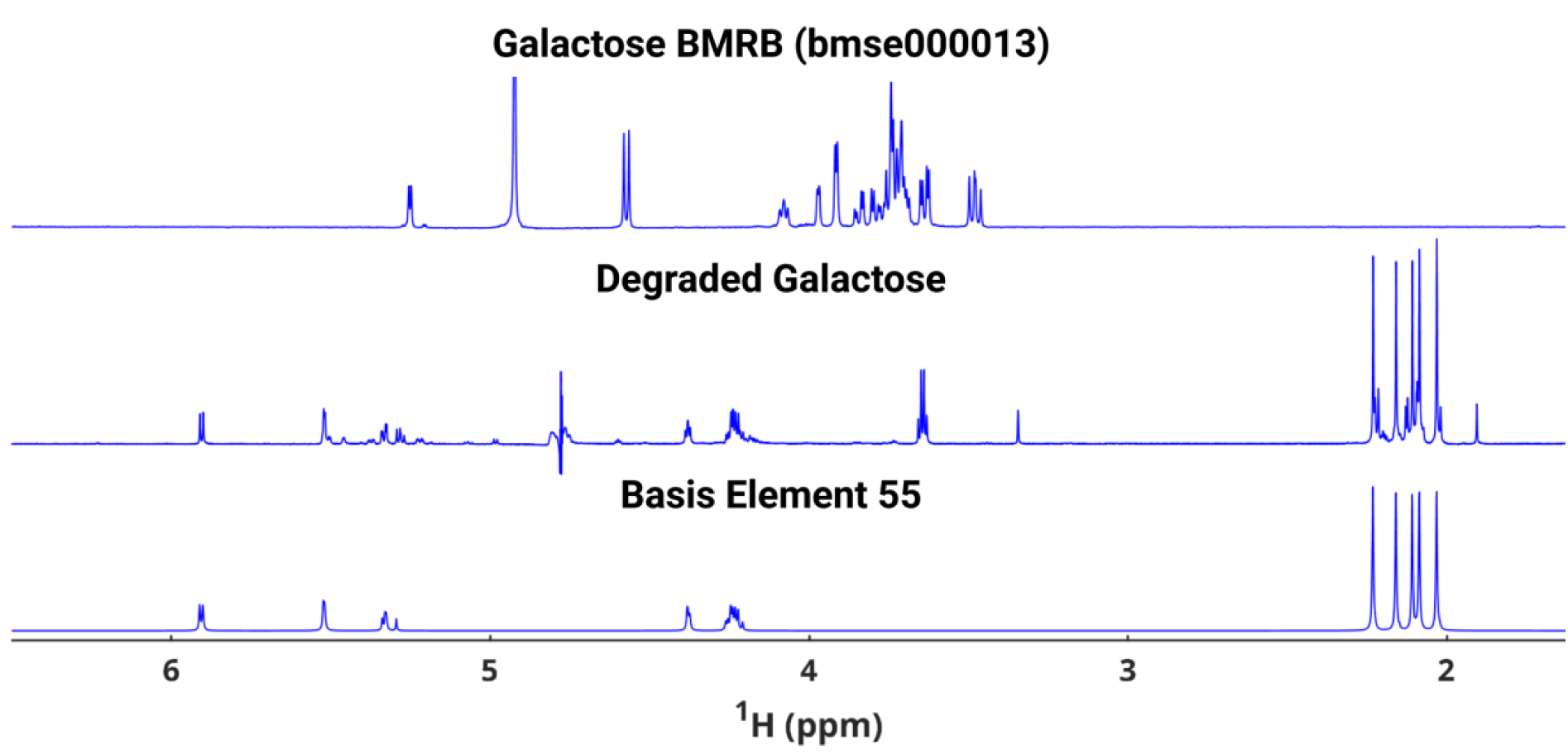
BMRB reference spectrum (bmse000013) of galactose, the ground-truth spectrum of the degraded galactose sample, and metabolite basis element 55. Unexpectedly, the galactose obtained for this study does not match the known spectrum of galactose from the BMRB database. The peaks in the galactose sample that are not captured in basis element 55 may be peaks of another contaminant.

**Figure Supplementary 5:**
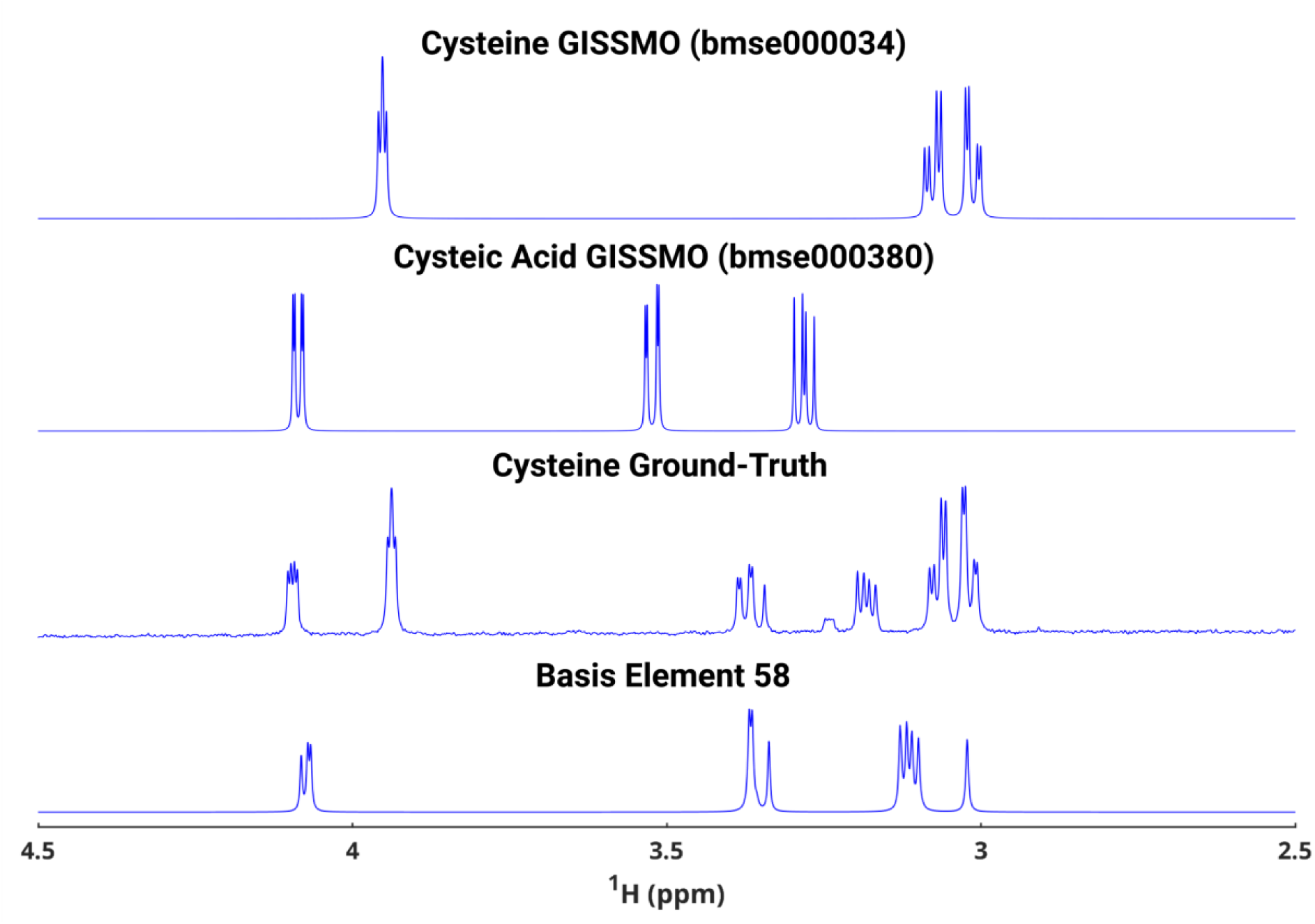
GISSMO reference spectrum (bmse000034) of cysteine, GISSMO reference spectrum (bmse000380) of cysteic acid, the ground-truth spectrum of the cysteine sample, and metabolite basis element 58. The cysteine ground-truth sample exhibits resonances that resemble those of cysteine and its oxidation product, cysteic acid. Changes in chemical shift are likely due to variations in pH. We quantified cysteic acid resonances in basis element 58.

**Figure Supplementary 6:**
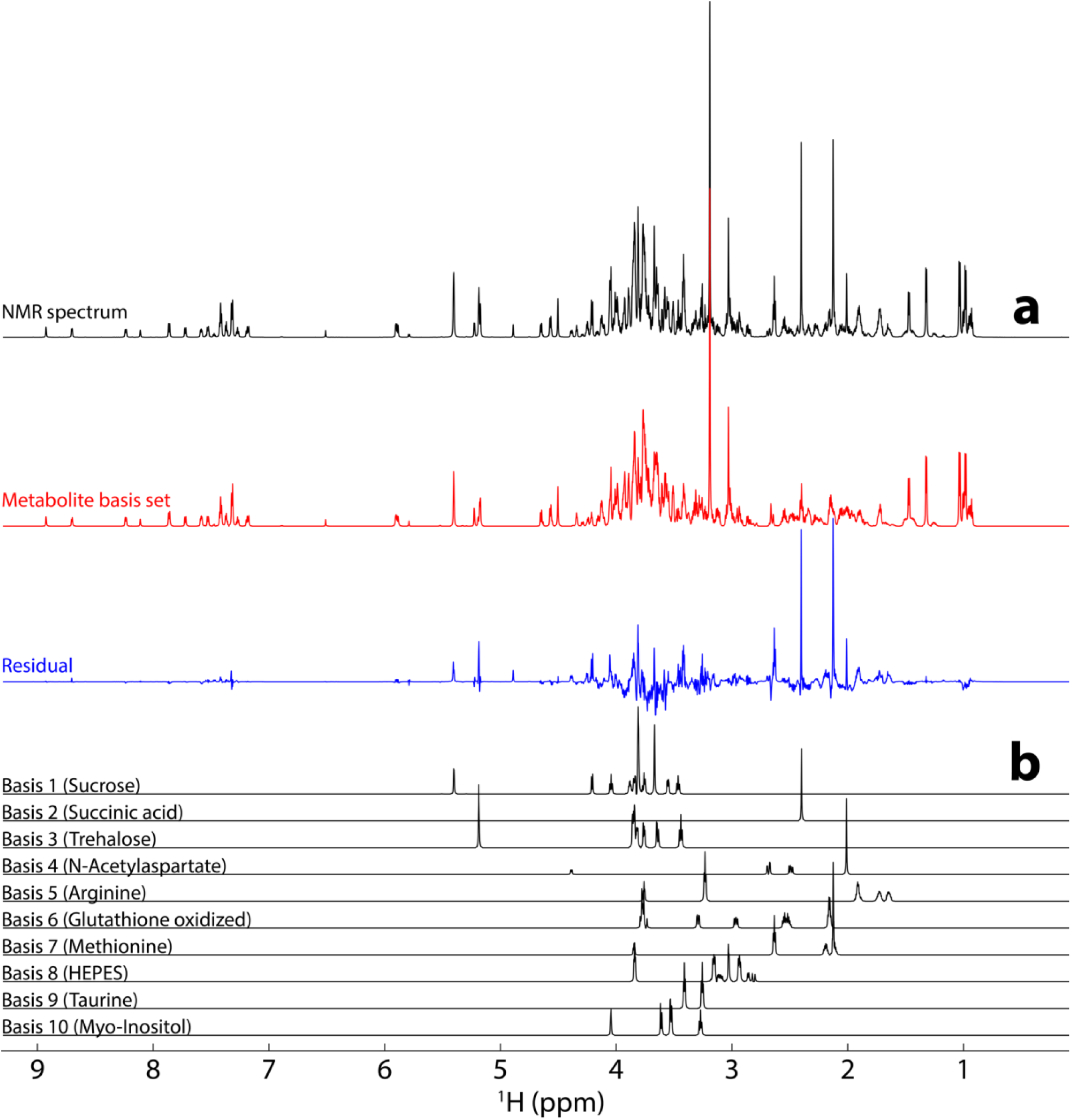
**a,** BATMAN fit of the mBS from our ground-truth set of 53 synthetic metabolite solutions with the first 10 mBS elements excluded from fitting. The NMR spectrum at the top (black) is one of ten ground-truth experimental mixtures for this study, and the same spectrum is shown in Figure 5. The metabolite basis set (mBS) shown in red is the BATMAN fit. The blue trace shows the residuals (wavelet fit) from the Bayesian analysis. **b,** Spectra of the first 10 basis set elements excluded from the fit. In this example, the quantification model accounts for 82% of the total spectral intensity.

**Figure Supplementary 7:**
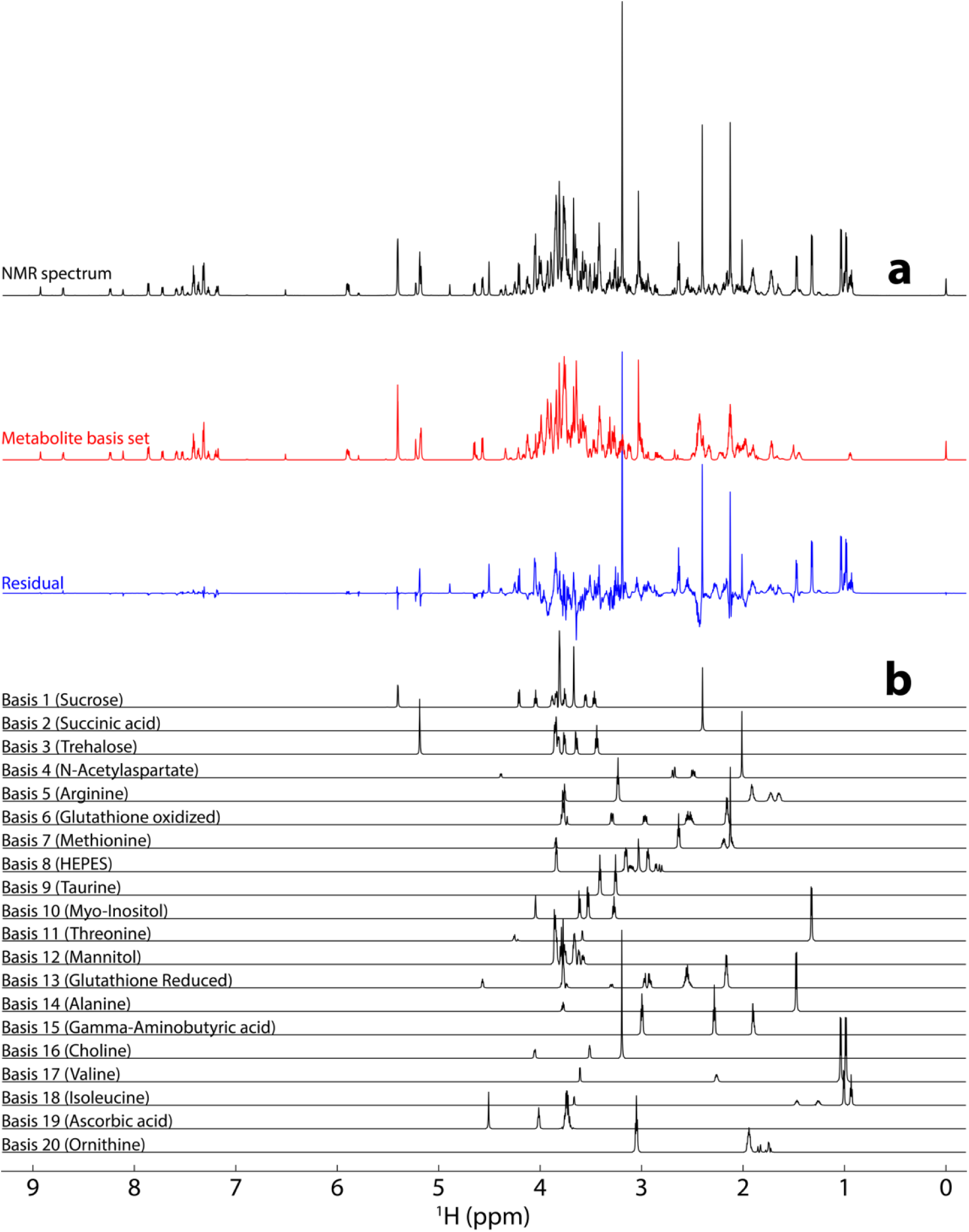
**a,** BATMAN fit of the mBS from our ground-truth set of 53 synthetic metabolite solutions with the first 20 mBS elements excluded from fitting. The NMR spectrum at the top (black) is one of ten ground-truth experimental mixtures for this study, and the same spectrum is shown in Figure 5. The metabolite basis set (mBS) shown in red is the BATMAN fit. The blue trace shows the residuals (wavelet fit) from the Bayesian analysis. **b,** Spectra of the first 20 basis set elements excluded from the fit. In this example, the quantification model accounts for 56% of the total spectral intensity.

**Figure Supplementary 8:**
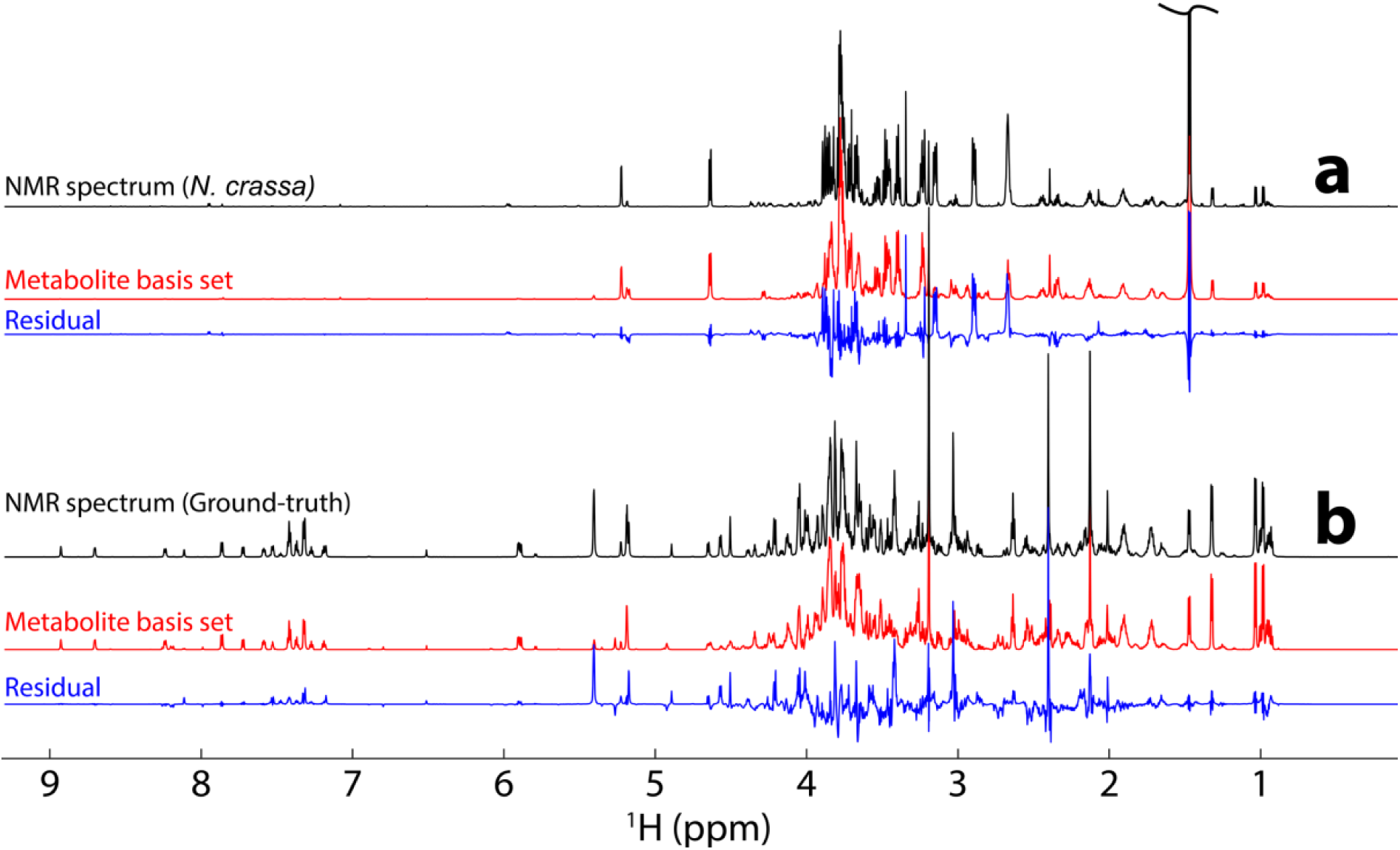
**a,** BATMAN fit of the mBS from the ground-truth mFL to the *N. crassa* unfractionated mixture spectrum. The BATMAN fit explained 83% of the original spectral intensity. **b,** BATMAN fit of the mBS from the *N. crassa* mFL to one of the ground-truth mixture spectra. In this example, the quantification model accounts for 82% of the total spectral intensity.

### Ground-Truth Methods

#### Creating Ground-Truth Spectra

The compounds used to create the single-component ground-truth reference spectra were common metabolites readily available in the laboratory. The milligram amount required for all 53 metabolites was calculated to prepare a 5 mL solution at a 100 mM concentration in H_2_O. An approximate amount of powder was weighed for each metabolite and added to a 10 mL glass centrifugal tube. LC/MS grade H_2_O was added, the tubes were vortexed, and stored at -20 °C. Due to solubility issues, not all metabolites fully dissolved in the solution.

#### Preparing Fractionation Mixture

10 µL was taken from each single-metabolite reference sample except for aspartate, in which 20 µL was taken and added to a 2 mL screw cap centrifuge tube. The tube was dried in a Centrivap and resuspended in 400 µL of 80:20 LC/MS grade MeOH: H_2_O.

#### HPLC Fractionation

The fraction library was produced by three 100 µL injections using an Agilent 1260 Infinity HPLC with XBridge BEH Amide OBD Prep Column, 130 Å, 5 µm, 10 x 250 mm HILIC column at 25°C. Full scan data was collected using an Agilent Infinity Lab Single Quadrupole MSD in positive ion mode (50-1250 Da). OpenLab ChemStation software was used for data acquisition and visualization.

A 38 min linear gradient of 0.1% formic acid in H_2_O (A) and 0.1% formic acid in ACN (B) was used for the fractionation. From 0-20 min, a linear gradient of 5% to 30% A was used, followed by a linear gradient of 30% to 50% A from 20-30 min, all at a flow rate of 3.5 mL/min. From 30-35 min, a linear gradient of 50% to 65% A was used, followed by an isocratic hold from 35-38 min, both at a flow rate of 2 mL/min. A post-time of 8 min was set to allow the system to equilibrate to the initial condition of 5% A before further injections. Between 0.9 and 30 min, 140 equally spaced fractions were collected, approximately 12.5 s per fraction. Fractionation was done over identical fraction vials for all three injections. Between injections 2 and 3 and after injection 3, the vials were dried using a Centrivap. The dried fraction vials were stored at -80 °C before NMR data collection.

#### Creation of Mixtures for Fitting

Ten mixtures of all metabolites were created. A random number generator was used to generate values between 1-55 µL to decide how much of each single-compound ground-truth sample would be added to each mixture (Table Supplementary 4). The mixtures were then dried using a Centrivap and reconstituted in 550 µL of buffered D_2_O (100 mM sodium phosphate buffer and 0.333 mM DSS-D_6_ at 7.4 pH). 55 µL of the solutions were pipetted into 1.7 mm Bruker SampleJet NMR tubes.

#### NMR Data Acquisition

The fractions were reconstituted in 55 μL D_2_O buffer and transferred to 1.7 mm Bruker SampleJet tubes. Solvent blanks were placed in positions 1, 96, 97, and 144 amongst the fractions. NMR data were collected using a Bruker Avance Neo console on an Oxford 800 MHz magnet with a 1.7 mm TCI cryoprobe and a cooled SampleJet sample changer. One-dimensional NMR data were acquired at 298K using a “noesypr1d” pulse sequence, and 32,768 points were collected with 8 dummy scans and 64 scans for each sample. The ten mixtures were run on the same instrument. The data were acquired at 298K using a “noesypr1d” pulse sequence, and 32,768 points were collected with 4 dummy scans and 16 scans for each sample. The data were automatically updated to NAN and NMRbox for processing.

#### Data processing and SAND

Prior to applying SAND, the spectra for ground-truth reference samples and mixtures were processed using the NMRPipe batch processing scheme, and reference deconvolution was applied using DSS as the reference lineshape, with a target linewidth of 1 Hz and a line broadening of 1.5 Hz. The reference samples were then time-domain modeled by SAND over the range of 9.1 ppm to -0.15 ppm.

#### Relative Concentrations of Basis Set Elements

Because of the limited solubility of some compounds, the expected concentrations do not always match the actual concentrations of the metabolites in the mixtures. To account for this, metabolite concentrations in the mixtures were determined by comparing the integral of isolated resonances to the integral of DSS using Mnova software. The newly calculated concentrations and the BATMAN relative concentrations were then compared in MATLAB.

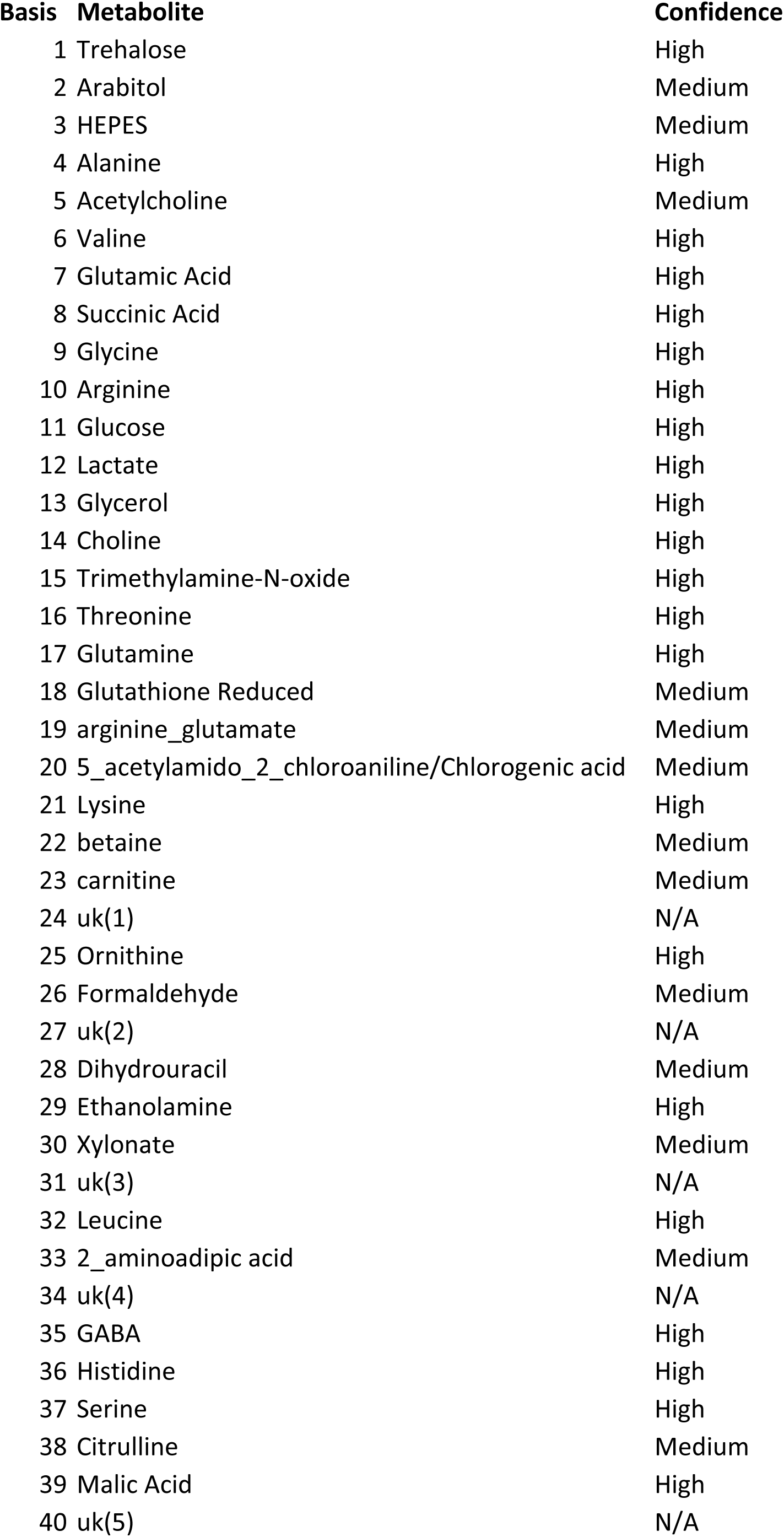
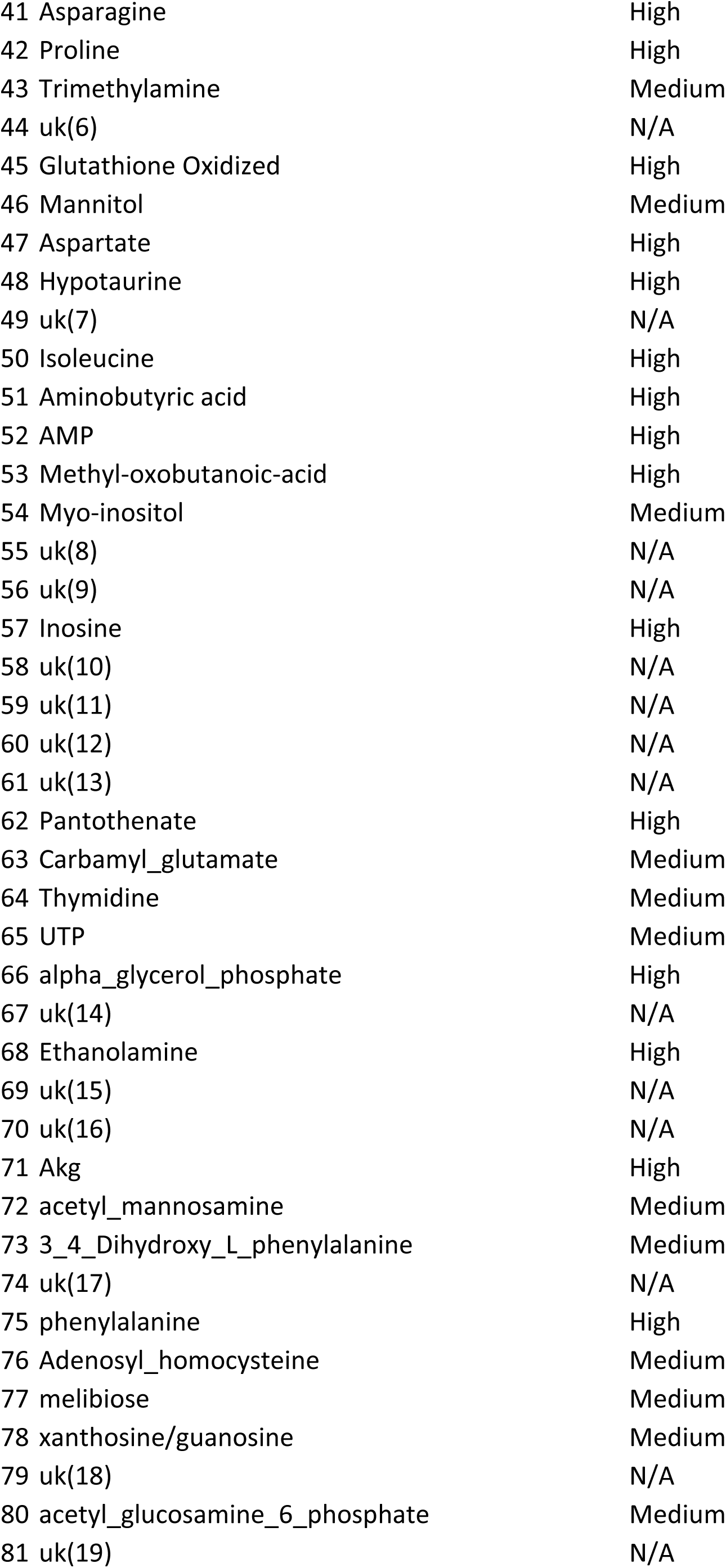
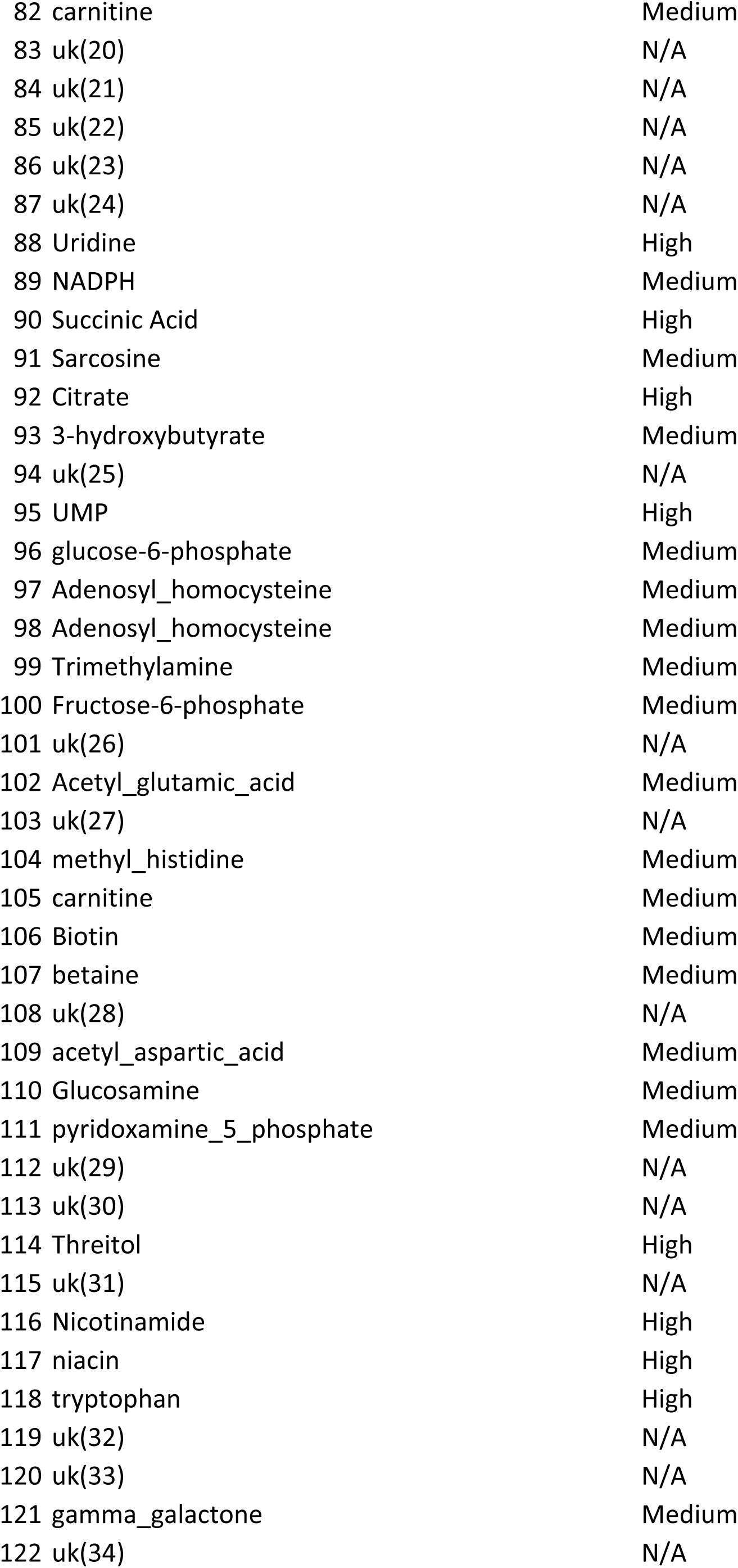
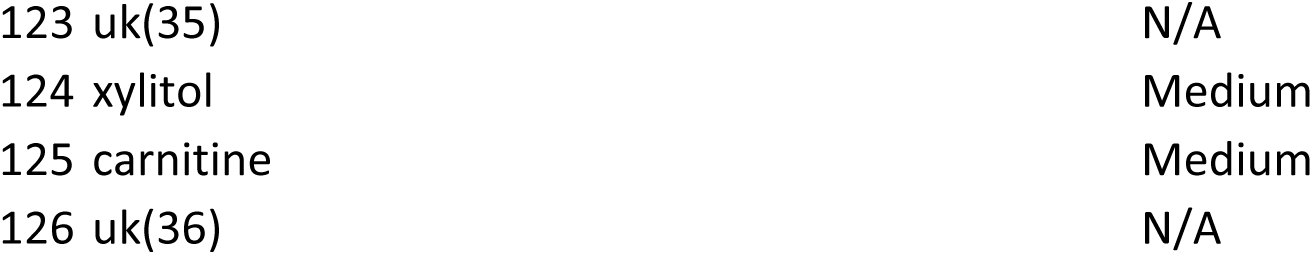

| **Metabolite** | **Basis** |
| --- | --- |
| 2-aminoadipic acid | Basis 46 |
| Adenine | Basis 60 |
| Alanine | Basis 14 |
| AKG | Basis 41 |
| Arginine | Basis 5 |
| Ascorbic acid | Basis 19 |
| Asparagine | Basis 39 |
| Aspartate | Basis 37 |
| Benzoic acid | Basis 59 |
| Choline | Basis 16 |
| Creatinine | Basis 25 |
| Cysteine | Basis 58* |
| Fructose | Basis 38 |
| Fumaric acid | Basis 47 |
| GABA | Basis 15 |
| Galactose | Basis 55* |
| Glucose | Basis 36 |
| Glutamic Acid | Basis 40 |
| Glutamine | Basis 30 |
| Glutathione Oxidized | Basis 6 |
| Glutathione Reduced | Basis 13 |
| Glycine | Basis 22 |
| HEPES | Basis 8 |
| Histidine | Basis 29 |
| Isoleucine | Basis 18 |
| Leucine | Basis 26 |
| Lysine | Basis 21 + Basis 34 |
| Malic Acid | Basis 33 |
| Maltose | Basis 23 |
| Mannitol | Basis 12 |
| Mannose | Basis 35 |
| Methionine | Basis 7 |
| myo-Inositol | Basis 10 |
| N-acetyl Aspartic acid | Basis 4 |
| Nicotinamide | Basis 45 |
| Ornithine | Basis 20 + Basis 32 |
| Phenylalanine | Basis 28 |
| Proline | Basis 42 |
| Pyruvate |  |
| Quinic Acid | Basis 24 |
| Serine | Basis 27 |
| Sorbitol | Basis 49 |
| Succinic acid | Basis 2 |
| Sucrose | Basis 1 |
| Taurine | Basis 9 |
| Threonine | Basis 11 |
| Trehalose | Basis 3 |
| Tryptophan | Basis 44 |
| Tyrosine | Basis 57 |
| Uracil | Basis 56 |
| Uridine | Basis 31 |
| Valine | Basis 17 |
| Xylose | Basis 43 |

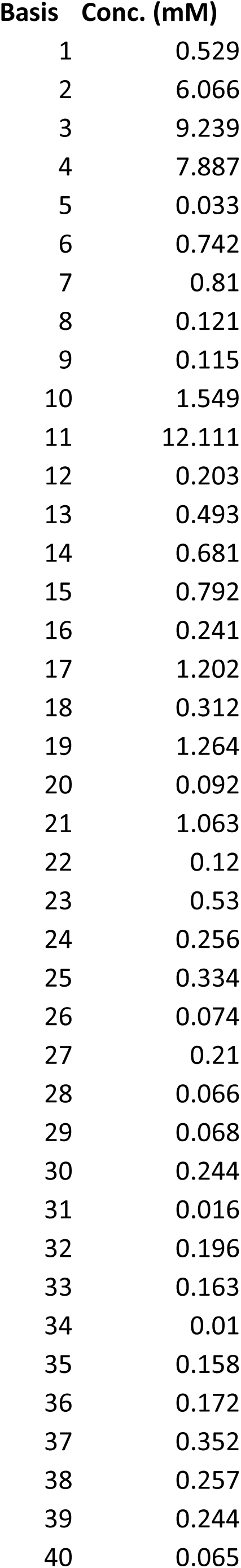
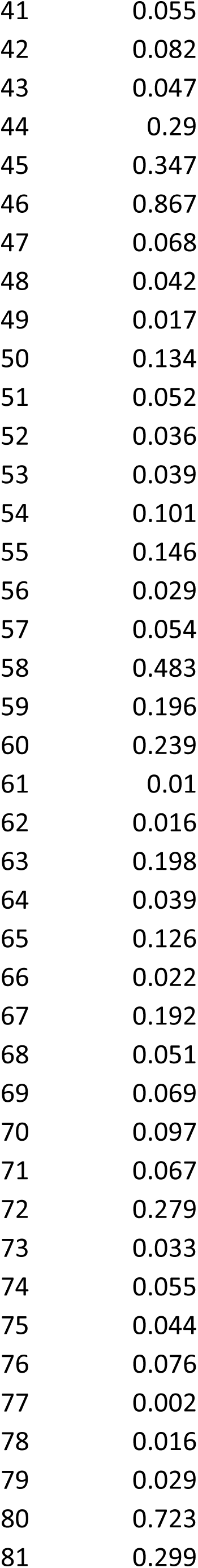
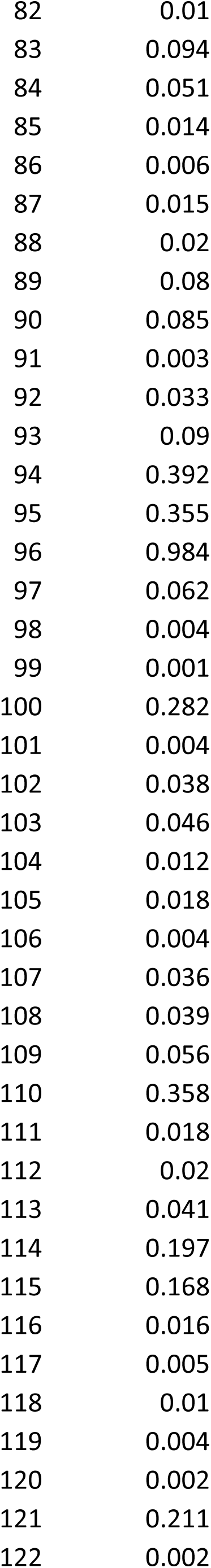
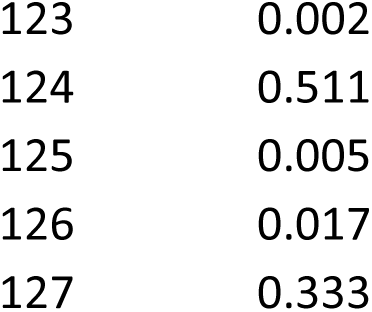

|  | Mixture 1 | Mixture 2 | Mixture 3 | Mixture 4 | Mixture 5 | Mixture 6 |
| --- | --- | --- | --- | --- | --- | --- |
| Ornithine | 51 | 51 | 2 | 21 | 41 | 46 |
| Asparagine | 7 | 39 | 5 | 43 | 25 | 50 |
| Glycine | 54 | 37 | 15 | 3 | 28 | 6 |
| Choline | 18 | 18 | 5 | 44 | 9 | 30 |
| Histidine | 37 | 33 | 19 | 14 | 3 | 14 |
| Taurine | 28 | 17 | 14 | 50 | 12 | 2 |
| Maltose | 48 | 43 | 34 | 54 | 35 | 2 |
| Fumaric acid | 5 | 29 | 34 | 52 | 45 | 48 |
| Glutathione Oxidized | 4 | 20 | 31 | 34 | 15 | 34 |
| Sorbitol | 43 | 12 | 20 | 31 | 4 | 4 |
| Uridine | 26 | 6 | 45 | 20 | 27 | 46 |
| Valine | 52 | 36 | 45 | 40 | 12 | 45 |
| Adenine | 21 | 32 | 4 | 10 | 4 | 39 |
| Serine | 15 | 22 | 19 | 18 | 50 | 20 |
| 2-aminoadipic acid | 20 | 14 | 13 | 26 | 27 | 43 |
| Quinic acid | 3 | 25 | 24 | 12 | 4 | 31 |
| Leucine | 15 | 35 | 4 | 11 | 54 | 13 |
| Malic acid | 35 | 42 | 28 | N/A | 26 | 24 |
| Creatinine | 46 | 48 | 41 | 37 | 30 | 1 |
| Proline | 3 | 37 | 3 | 32 | 21 | 42 |
| Sucrose | 15 | 28 | 17 | 53 | 25 | 8 |
| Methionine | 43 | 47 | 10 | 55 | 54 | 44 |
| Glutamic acid | 43 | 2 | 39 | 17 | 18 | 44 |
| Glucose | 19 | 14 | 40 | 4 | 49 | 50 |
| Lysine | 13 | 19 | 31 | 39 | 28 | 12 |
| HEPES | 13 | 54 | 6 | 23 | 50 | 6 |
| Ascorbic acid | 6 | 43 | 34 | 47 | 31 | 7 |
| Alanine | 1 | 25 | 42 | 22 | 55 | 23 |
| Isoleucine | 43 | 41 | 12 | 16 | 54 | 25 |
| AKG | 29 | 37 | 4 | 23 | 48 | 24 |
| Arginine | 47 | 52 | 23 | 45 | 21 | 28 |
| Trehalose | 48 | 25 | 48 | 31 | 54 | 15 |
| Xylose | 13 | 19 | 16 | 39 | 32 | 52 |
| Aspartate | 22 | 46 | 29 | 2 | 55 | 10 |
| Uracil | 44 | 18 | 23 | 51 | 27 | 24 |
| N-acetyl Aspartic acid | 24 | 30 | 52 | 19 | 6 | 46 |
| GABA | 39 | 17 | 17 | 7 | 47 | 10 |
| Myo-Inositol | 3 | 46 | 20 | 7 | 48 | 31 |
| Tyrosine | 54 | 40 | 39 | 29 | 14 | 36 |
| Pyruvate | 30 | 2 | 14 | 19 | 54 | 23 |
| Cysteine | 32 | 15 | 34 | 24 | 26 | 46 |
| Succinic acid | 12 | 49 | 35 | 45 | 5 | 18 |
| Nicotinamide | 43 | 23 | 45 | 21 | 33 | 55 |
| Tryptophan | 35 | 31 | 47 | 40 | 31 | 7 |
| Threonine | 21 | 22 | 15 | 43 | 54 | 30 |
| Benzoic acid | 18 | 43 | 6 | 30 | 33 | 9 |
| Glutathione Reduced | 54 | 29 | 17 | 48 | 50 | 6 |
| Fructose | 18 | 31 | 55 | 26 | 31 | 22 |
| Galactose | 39 | 4 | 37 | 4 | 35 | 51 |
| Glutamine | 20 | 26 | 42 | 9 | 35 | 46 |
| Mannitol | 4 | 53 | 53 | 28 | 18 | 42 |
| Mannose | 3 | 29 | 52 | 54 | 23 | 37 |
| Phenylalanine | 8 | 28 | 21 | 35 | 37 | 11 |

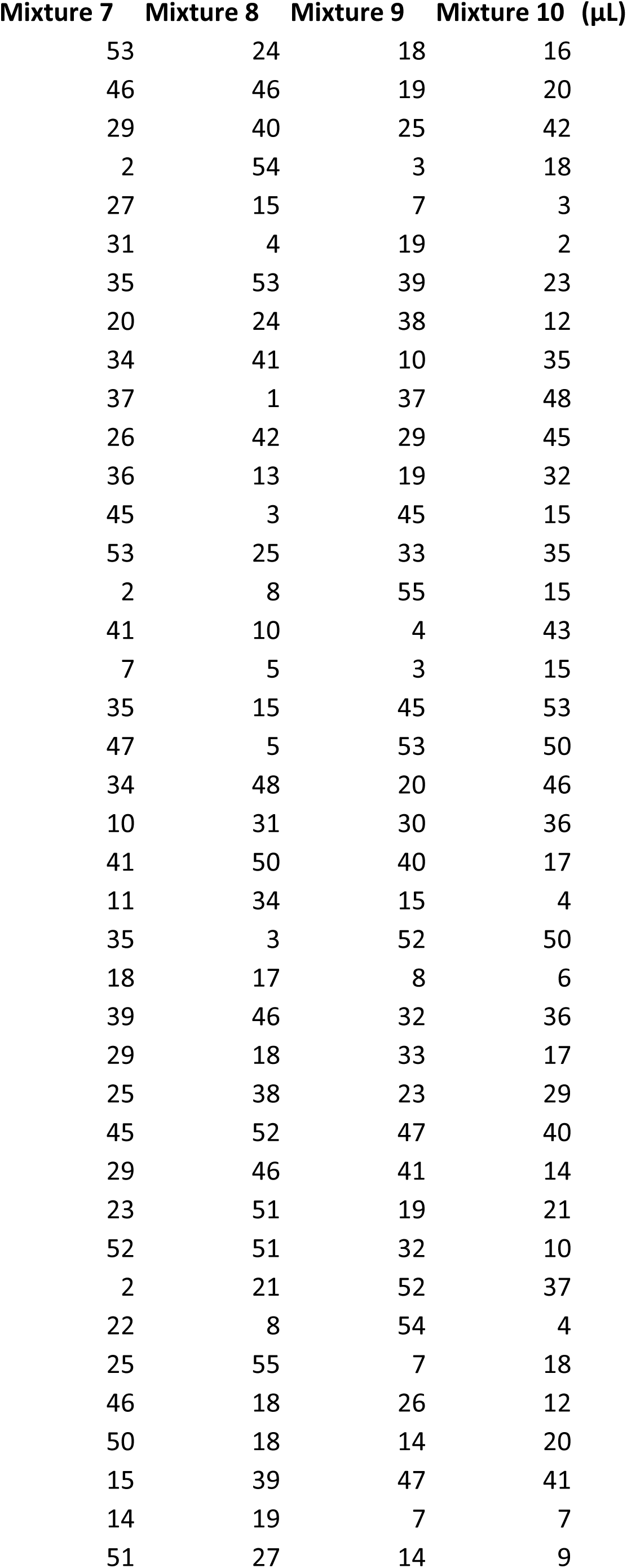
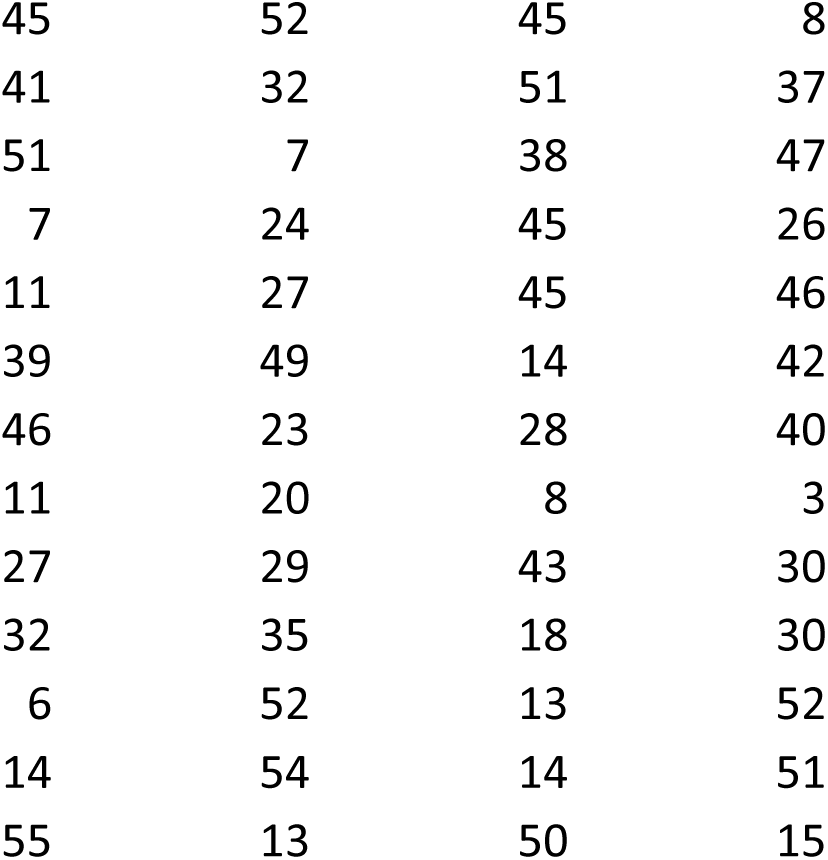

## References

1. Edison, A.S. et al. NMR: Unique Strengths That Enhance Modern Metabolomics Research. Anal Chem 93, 478–499 (2021).

2. Wang, M. et al. Sharing and community curation of mass spectrometry data with Global Natural Products Social Molecular Networking. Nat Biotechnol 34, 828–837 (2016).

3. Blazenovic, I., Kind, T., Ji, J. & Fiehn, O. Software Tools and Approaches for Compound Identification of LC-MS/MS Data in Metabolomics. Metabolites 8 (2018).

4. Wishart, D.S. et al. NMR and Metabolomics-A Roadmap for the Future. Metabolites 12 (2022).

5. Edison, A.S. & Schroeder, F.C. (ed. L.L.H.W. Mander) 169–196 (Elsevier, Oxford; 2010).

6. Lewis, I.A., Schommer, S.C. & Markley, J.L. rNMR: open source software for identifying and quantifying metabolites in NMR spectra. Magnetic Resonance in Chemistry 47 **Suppl 1**, S123–126 (2009).

7. Bingol, K., Li, D.W., Zhang, B. & Bruschweiler, R. Comprehensive Metabolite Identification Strategy Using Multiple Two-Dimensional NMR Spectra of a Complex Mixture Implemented in the COLMARm Web Server. Anal Chem 88, 12411–12418 (2016).

8. Bingol, K. & Bruschweiler, R. Multidimensional approaches to NMR-based metabolomics. Anal Chem 86, 47–57 (2014).

9. Uchimiya, M. et al. (13)C NMR metabolomics: J-resolved STOCSY meets INADEQUATE. J Magn Reson 347, 107365 (2023).

10. Blaise, B.J. et al. Statistical analysis in metabolic phenotyping. Nat Protoc (2021).

11. Robinette, S.L., Lindon, J.C. & Nicholson, J.K. Statistical spectroscopic tools for biomarker discovery and systems medicine. Analytical Chemistry 85, 5297–5303 (2013).

12. Cloarec, O. et al. Statistical total correlation spectroscopy: an exploratory approach for latent biomarker identification from metabolic 1H NMR data sets. Analytical Chemistry 77, 1282–1289 (2005).

13. Wei, S. et al. Ratio analysis nuclear magnetic resonance spectroscopy for selective metabolite identification in complex samples. Analytical Chemistry 83, 7616–7623 (2011).

14. Clendinen, C.S., Pasquel, C., Ajredini, R. & Edison, A.S. (13)C NMR Metabolomics: INADEQUATE Network Analysis. Anal Chem 87, 5698–5706 (2015).

15. Metz, T.O. et al. Introducing “Identification Probability” for Automated and Transferable Assessment of Metabolite Identification Confidence in Metabolomics and Related Studies. Anal Chem (2024).

16. Mercier, P., Lewis, M.J., Chang, D., Baker, D. & Wishart, D.S. Towards automatic metabolomic profiling of high-resolution one-dimensional proton NMR spectra. Journal of Biomolecular NMR 49, 307–323 (2011).

17. Wishart, D.S. Human Metabolome Database: completing the ‘human parts list’. Pharmacogenomics 8, 683–686 (2007).

18. Tredwell, G.D., Behrends, V., Geier, F.M., Liebeke, M. & Bundy, J.G. Between-person comparison of metabolite fitting for NMR-based quantitative metabolomics. Analytical Chemistry 83, 8683–8687 (2011).

19. Loo, R.L. et al. Quantitative In-Vitro Diagnostic NMR Spectroscopy for Lipoprotein and Metabolite Measurements in Plasma and Serum: Recommendations for Analytical Artifact Minimization with Special Reference to COVID-19/SARS-CoV-2 Samples. J Proteome Res 19, 4428–4441 (2020).

20. Dona, A.C. et al. Precision high-throughput proton NMR spectroscopy of human urine, serum, and plasma for large-scale metabolic phenotyping. Analytical Chemistry 86, 9887–9894 (2014).

21. Ravanbakhsh, S. et al. Accurate, fully-automated NMR spectral profiling for metabolomics. PLoS One 10, e0124219 (2015).

22. Hao, J. et al. Bayesian deconvolution and quantification of metabolites in complex 1D NMR spectra using BATMAN. Nat Protoc 9, 1416–1427 (2014).

23. Gouveia, G.J. et al. Long-Term Metabolomics Reference Material. Anal Chem 93, 9193–9199 (2021).

24. Wu, Y. et al. SAND: Automated Time-Domain Modeling of NMR Spectra Applied to Metabolite Quantification. Anal Chem 96, 1843–1851 (2024).

25. Robinette, S.L., Bruschweiler, R., Schroeder, F.C. & Edison, A.S. NMR in metabolomics and natural products research: two sides of the same coin. Acc Chem Res 45, 288–297 (2012).

26. Sumner, L.W., Lei, Z., Nikolau, B.J. & Saito, K. Modern plant metabolomics: advanced natural product gene discoveries, improved technologies, and future prospects. Nat Prod Rep 32, 212–229 (2015).

27. Wolfender, J.L., Nuzillard, J.M., van der Hooft, J.J.J., Renault, J.H. & Bertrand, S. Accelerating metabolite identification in natural product research: toward an ideal combination of LC-HRMS/MS and NMR profiling, in silico databases and chemometrics. Anal Chem (2018).

28. Appiah-Amponsah, E., Owusu-Sarfo, K., Gowda, G.A., Ye, T. & Raftery, D. Combining Hydrophilic Interaction Chromatography (HILIC) and Isotope Tagging for Off-Line LC-NMR Applications in Metabolite Analysis. Metabolites 3, 575–591 (2013).

29. Whiley, L. et al. Systematic Isolation and Structure Elucidation of Urinary Metabolites Optimized for the Analytical-Scale Molecular Profiling Laboratory. Anal Chem 91, 8873–8882 (2019).

30. Krishnamurthy, K. CRAFT (complete reduction to amplitude frequency table) - robust and time-efficient Bayesian approach for quantitative mixture analysis by NMR. Magnetic Resonance in Chemistry 51, 821–829 (2013).

31. Li, D.W., Bruschweiler-Li, L., Hansen, A.L. & Bruschweiler, R. DEEP Picker1D and Voigt Fitter1D: a versatile tool set for the automated quantitative spectral deconvolution of complex 1D-NMR spectra. Magn Reson (Gott) 4, 19–26 (2023).

32. Li, D.-W., Hansen, A.L., Yuan, C., Bruschweiler-Li, L. & Brüschweiler, R. DEEP picker is a deep neural network for accurate deconvolution of complex two-dimensional NMR spectra. Nature Communications 12 (2021).

33. Dashti, H. et al. Spin System Modeling of Nuclear Magnetic Resonance Spectra for Applications in Metabolomics and Small Molecule Screening. Anal Chem 89, 12201–12208 (2017).

34. Nielsen, N.-P.V., Carstensen, J.M. & Smedsgaard, J. Aligning of single and multiple wavelength chromatographic profiles for chemometric data analysis using correlation optimised warping. Journal of Chromatography A 805, 17–35 (1998).

35. Wong, J.W.H., Durante, C. & Cartwright, H.M. Application of Fast Fourier Transform Cross-Correlation for the Alignment of Large Chromatographic and Spectral Datasets. Analytical Chemistry 77, 5655–5661 (2005).

36. Wu, W. et al. Peak Alignment of Urine NMR Spectra Using Fuzzy Warping. Journal of Chemical Information and Modeling 46, 863–875 (2006).

37. Savorani, F., Tomasi, G. & Engelsen, S.B. icoshift: A versatile tool for the rapid alignment of 1D NMR spectra. Journal of Magnetic Resonance 202, 190–202 (2010).

38. Metzenberg, R.L. in Fungal Genetics Reports, Vol. 51 (2004).

39. Maciejewski, M.W. et al. NMRbox: A Resource for Biomolecular NMR Computation. Biophys J 112, 1529–1534 (2017).

40. Metz, K.R., Lam, M.M. & Webb, A.G. Reference deconvolution: A simple and effective method for resolution enhancement in nuclear magnetic resonance spectroscopy. Concepts in Magnetic Resonance 12, 21–42 (2000).

41. Morris, G.A., Barjat, H. & Home, T.J. Reference deconvolution methods. Progress in Nuclear Magnetic Resonance Spectroscopy 31, 197–257 (1997).

42. Barjat, H., Morris, G.A., Swanson, A.G., Smart, S. & Williams, S.C.R. Reference Deconvolution Using Multiplet Reference Signals. Journal of Magnetic Resonance, Series A 116, 206–214 (1995).

43. Hu, H., Van, Q.N., Mandelshtam, V.A. & Shaka, A.J. Reference deconvolution, phase correction, and line listing of NMR spectra by the 1D filter diagonalization method. Journal of Magnetic Resonance 134, 76–87 (1998).

44. Hu, H., Van, Q.N., Mandelshtam, V.A. & Shaka, A.J. Reference deconvolution, phase correction, and line listing of NMR spectra by the 1D filter diagonalization method. J Magn Reson 134, 76–87 (1998).

45. Li, D.W. et al. COLMAR1d: A Web Server for Automated, Quantitative One-Dimensional Nuclear Magnetic Resonance-Based Metabolomics at Arbitrary Magnetic Fields. Anal Chem 96, 17174–17183 (2024).

46. Doreleijers, J.F. et al. BioMagResBank database with sets of experimental NMR constraints corresponding to the structures of over 1400 biomolecules deposited in the Protein Data Bank. Journal of Biomolecular NMR 26, 139–146 (2003).

